# Bayesian Nonparametric Inference of Population Size Changes from Sequential Genealogies

**DOI:** 10.1101/019216

**Authors:** Julia A. Palacios, John Wakeley, Sohini Ramachandran

**Affiliations:** Department of Organismic and Evolutionary Biology, Harvard University; Department of Ecology and Evolutionary Biology, Brown University; Center for Computational Molecular Biology, Brown University

**Keywords:** Markov Process, Genomics, Sequentially Markov Coalescent, Point Process, Gaussian Process

## Abstract

Sophisticated inferential tools coupled with the coalescent model have recently emerged for estimating past population sizes from genomic data. Accurate methods are available for data from a single locus or from independent loci. Recent methods that model recombination require small sample sizes, make constraining assumptions about population size changes, and do not report measures of uncertainty for estimates. Here, we develop a Gaussian process-based Bayesian nonparametric method coupled with a sequentially Markov coalescent model which allows accurate inference of population sizes over time from a set of genealogies. In contrast to current methods, our approach considers a broad class of recombination events, including those that do not change local genealogies. We show that our method outperforms recent likelihood-based methods that rely on discretization of the parameter space. We illustrate the application of our method to multiple demographic histories, including population bottlenecks and exponential growth. In simulation, our Bayesian approach produces point estimates four times more accurate than maximum likelihood estimation (based on the sum of absolute differences between the truth and the estimated values). Further, our method’s credible intervals for population size as a function of time cover 90 percent of true values across multiple demographic scenarios, enabling formal hypothesis testing about population size differences over time. Using genealogies estimated with *ARGweaver*, we apply our method to European and Yoruban samples from the 1000 Genomes Project and confirm key known aspects of population size history over the past 150,000 years.

## 1 Introduction

For a single non-recombining locus, neutral coalescent theory predicts the set of timed ancestral relationships among sampled individuals, known as a gene genealogy (Kingman 1982; Hudson 1983; Tajima 1983; Hudson 1990). In the coalescent model with variable population size, the rate at which two lineages coalesce, or have a common ancestor, is a function of the population size in the past. Here we denote the *population size trajectory* by *N*(*t*), where *t* is time in the past, and use the term *local genealogy* to describe ancestral relationships at one non-recombining locus. When analyzing multilocus sequences, a single local genealogy will not represent the full history of the sample. Instead, the set of ancestral relationships and recombination events among a sample of multilocus sequences can be represented by a graph, known as the ancestral recombination graph (ARG) which depicts the complex structure of neighboring local genealogies and results in a computationally expensive model for inferring *N*(*t*) (Griffiths and Marjoram 1997; Wiuf and Hein 1999).

Recent studies have leveraged computationally simpler approximations for the coalescent with recombination—the sequentially Markov coalescent (SMC) (McVean and Cardin 2005) and its variant SMC′ (Marjoram and Wall 2006; Chen et al. 2009)—both of which model local genealogies as a continuous time Markov process along sequences (Figure 1). The difference between the SMC and SMC′ is that the SMC models only the class of recombination events that alter local genealogies of the sample. In general, the SMC′ is a better approximation to the ARG than the SMC (Chen et al. 2009; Wilton et al. 2015). Because of these features, in this work we rely on the SMC′ to model local genealogies with recombination.

**Figure 1:**
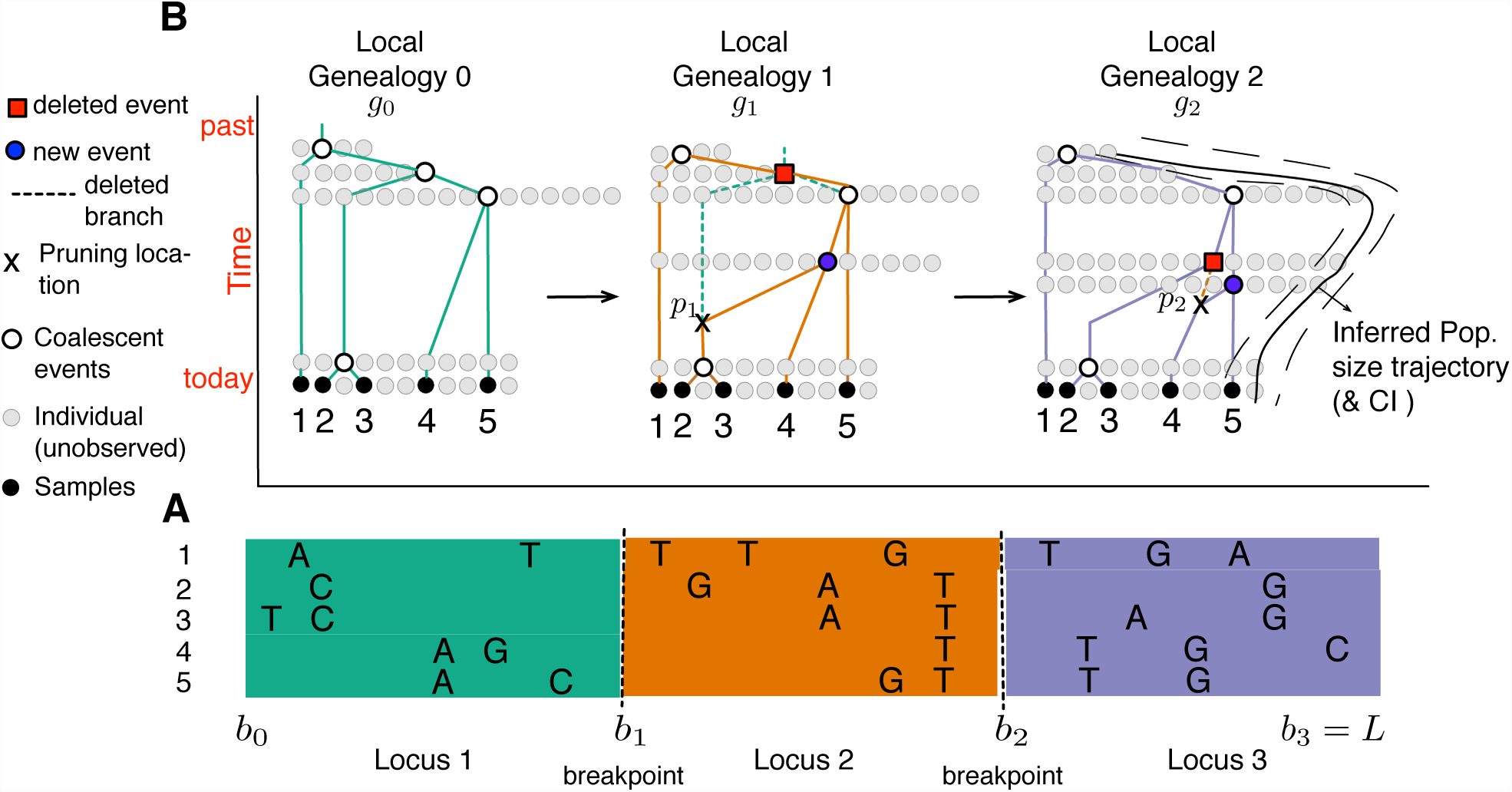
SMC′ model for inferring population size trajectories. Drawn after Rasmussen et al. (2014) to highlight notation specific to our study. **A.** Observed sequence data in a segment of length L from five individuals; three loci are shown delimited by recombination breakpoints *b*_1_ and *b*_2_. Only the derived mutations at polymorphic sites are shown. **B.** Corresponding local genealogies *g*_*i*_ for each locus i. The five sampled individuals are depicted as black filled circles. Local genealogies have a Markovian degree 1 dependency. Each inter-coalescent time (the time interval between coalescent events denoted as empty circles) provides information about past population size (number of gray filled circles at a given time point). Moving from left to right after recombination breakpoint *b*_1_, the pruning location *p*_1_ is selected from genealogy *g*_0_ and the pruned branch is regrafted back on the genealogy (blue filled circle). The coalescent event of *g*_0_ depicted as a red filled circle in *g*_1_ is deleted. This creates the next genealogy *g*_1_. The process continues until *L*. At *L*, the population size trajectory *N*(*t*) (depicted as a black curve superimposed on *g*_2_) can be inferred.

Under the coalescent and the sequentially Markov coalescent (SMC and SMC′) models, population size trajectories and sequence data are separated by two stochastic processes: *i*) *a state process* which describes the relationship between the population size trajectory and the set of local genealogies, and *ii*) *an observation process* which describes how the hidden local genealogies are observed through patterns of nucleotide diversity in the sequence data. The observation process includes mutation and genotyping error while the state process models coalescence. Sequence data are then used to make inferences of population size trajectories. In this paper, we restrict attention to the state process of local genealogies and show how inferences of population size trajectories can be made from them. We solve a number of key modeling and inference problems, and thus provide a basis for developing efficient algorithms to infer population parameters from sequence data directly.

Whole-genome inference of population size trajectories has been hampered by the enormous size of the state space of local genealogies when the sample size is large. The pioneering, pairwise sequentially Markov Coalescent (PSMC) method of Li and Durbin (2011) employed the SMC to make inferences from a sample of size two (n = 2). In this method, time is discretized and the population size trajectory is piece-wise constant, allowing pairwise genealogies also to be discretized. Subsequent methods for samples larger than two similarly rely on the discretization of time and genealogies. The natural extension of the PSMC to *n* > 2 is the multiple sequentially Markovian coalescent (MSMC) (Schiffels and Durbin 2014). However, the MSMC models only the most recent coalescent event of the sample, and hence its estimation of population sizes is limited to very recent times. Other recent methods propose efficient ways of exploring the state space of hidden genealogies for *n* > 2 (Sheehan et al. 2013; Rasmussen et al. 2014), yet also rely on discretizing the state space of local genealogies and assume a piece-wise constant trajectory of population sizes. We show that the *a priori* specification of change points for the piece-wise population size trajectory required by current approaches is problematic because estimates of *N*(*t*) are sensitive to this specification. Moreover, current methods do not generate interval estimates for *N*(*t*).

Gaussian Process-based Bayesian inference of population size trajectories has proven to be a powerful and flexible nonparametric approach when applied to a single local genealogy (Palacios and Minin 2013; Lan et al. 2015). The two main advantages of the GP-based approach are: (*i*) it does not require a specific functional form of the population size trajectory (such as constant or exponential growth) and (*ii*) it does not require an arbitrary specification of change points in a piece-wise constant or linear framework.

In this paper, we show the downstream effects of discretizing time, assuming a piecewise constant trajectory, and reporting only point estimates for past population sizes. We overcome previous limitations by introducing a Bayesian nonparametric approach with a Gaussian Process (GP) to model the population size trajectory as a continuous function of time. More specifically, we model the logarithm of the population size trajectory *a priori* as a Gaussian process (the log ensures our estimates are positive). As mentioned above, we assume that local gene genealogies are known. For our Bayesian model, we develop a Markov Chain Monte Carlo (MCMC) method to sample from the posterior distribution of population sizes over time. Our MCMC algorithm uses the recently developed algorithm Split Hamiltonian Monte Carlo (splitHMC) (Shahbaba et al. 2014; Lan et al. 2015). splitHMC updates all model parameters jointly and it can be extended to a full inferential framework that is directly applicable to sequence data. In order to compare our Bayesian GP-based estimation of population size trajectories with a piece-wise constant maximum likelihood-based estimation (e.g. Li and Durbin 2011; Sheehan et al. 2013; Schiffels and Durbin 2014), we implemented the Expectation-maximization (EM) algorithm within our framework and computed the observed Fisher information to obtain confidence intervals of the maximum likelihood estimates.

Lastly, we address a key problem for inference of population size trajectories under sequentially Markov coalescent models is the efficient computation of transition densities needed in the calculation of likelihoods. Here, we express the transition densities of local genealogies in terms of local ranked tree shapes (Tajima 1983) and coalescent times, and show that these quantities are statistically sufficient for inferring population size trajectories either from sequence data directly or from the set of local genealogies. The use of ranked tree shapes allows us to exploit the state process of local genealogies efficiently since the space of ranked tree shapes has a smaller cardinality than the space of labeled topologies (Sainudiin et al. 2014).

## Methods: SMC′ Calculations

Following notation similar to Rasmussen et al. (2014) (Table 1) a realization of the embedded SMC′ chain consists of a set of *m* local genealogies (*g*_0_, *g*_1_,…,*g*_*m-*1_), *m* − 1 recombination break-points at chromosomal locations (*b*_1_, *b*_2_,…,*b*_*m-*1_), and *m* − 1 pruning locations (*p*_1_, *p*_2_,…,*p*_*m-*1_), where *p*_*i*_ = (*u*_*i*_, *w*_*i*_) indicates the time of the recombination event *u*_*i*_ and the branch *w*_*i*_ where recombination happened in genealogy *g*_*i-*1_ (Figure 1). Genealogy *g*_0_ corresponds to the genealogy of *n* sequences that contains the set of timed ancestral relationships among the *n* individuals for the chromosomal segment (0, *b*_1_]. Genealogy *g*_*i*_ corresponds to the genealogy of the same *n* sequences for the chromosomal segment (*b*_*i*_, *b*_*i*+1_] for *i* = 1, 2,…,*m* − 2. Finally, 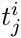 denotes the time when two of *j* lineages coalesce in genealogy *g*_*i*_, measured in units of generations before present.

**Table 1:** Notation for the SMC′ model used in this work.

|  | Symbol | Description |
| --- | --- | --- |
| Parameters: | $\rho$ | Recombination rate per site per generation |
| | $N(t)$ | Effective population size trajectory with time measured in units of $N_0$ generations |
| | $\tau$ | Hyperparameter that controls the smoothness of the log-Gaussian process prior on $N(t)$ |
| Notation specific to SMC' chain: | $L$ | Length of observed sequences |
| | $b_i$ | Chromosomal location of the $i$th recombination breakpoint |
| | $m$ | Number of local genealogies corresponding to $m - 1$ recombination events |
| | $s_{i+1} = b_{i+1} - b_i$ | Segment length for local genealogy $i$ |
| | $g_i$ | Local genealogy for the segment $(b_{i-1}, b_i)$ |
| Notation specific to local genealogy: | $n$ | Sample size, or number of sequences |
| | $l_i$ | Total tree length of local genealogy $g_i$ |
| | $A^i(t)$ | Piece-wise constant function of the number of ancestral lineages at time $t$ in local genealogy $g_i$ |
| | $t_j^i$ | Coalescent time in genealogy $g_i$ when two of $j$ lineages coalesce. $A^i(t_j^i -) = j$; $A^i(t_j^i +) = j - 1$ |
| | $\mathbf{t}^i = (t_n^i, t_{n-1}^i, \dots, t_2^i)$ | Vector of coalescent times of genealogy $g_i$ |
| | $p_i = (u_i, w_i)$ | Pruning location along local genealogy $g_i$ |
| | $u_i$ | Time when the recombination event happened along the height of the genealogy $g_i$ |
| | $w_i$ | Lineage on genealogy $g_{i-1}$ where the recombination event happened |
| | $w'_i$ | New lineage added on genealogy $g_i$ where the recombination event happened |
| | $t_{new}^i$ | Coalescent time in genealogy $g_i$ when the lineage $w_i$ coalesces. |
| | $t_{del}^i$ | Coalescent time in genealogy $g_{i-1}$ that no longer exists in genealogy $g_i$ |
| | $c_i$ | Lineage on genealogy $g_i$ that coalesces with lineage $w'_i$ |
| | $F_{j,k}^i$ | Number of free lineages in local genealogy $g_i$ that do not coalesce in the time interval $(t_{j+1}^i, t_k^i)$ |
| | $I^i(t)$ | Piece-wise constant function that takes values in $\{0, 1, 2\}$ indicating the number of ancestral lineages at time $t$ in genealogy $g_i$ where the pruning event would produce a visible transition to $g_{i+1}$ |
| Discretization: | $d$ | Number of change points at which $N(t)$ is estimated |
| | $\mathbf{x} = (x_1, \dots, x_d)$ | Times at which $N(t)$ is estimated |

Using capital letters to denote random variables, the evolution of the SMC′ process along chromosomal segments is governed by a point process *B* = {*B*_*i*_}_*i*∈ℕ_ that represents the random locations of recombination breakpoints. We use *S*_*i*_ = *B*_*i*_ − *B*_*i-*1_, for *i* = 1, 2,…, *m*, to denote the segment lengths for each local genealogy, with S_0_ = *B*_0_ = 0. Let *G* = {*G*_*i*_}_*i*∈ℕ_ be the chain which records the local genealogies, and let *P* = (*U, W*) = {(*U*_*i*_, *W*_*i*_)}_*i*∈ℕ_ represent the chain which records the pruning locations (time and branch) on G. The sequence (*G*_*i*_, *P*_*i*_ = {*U*_*i*_, *W*_*i*_}, *B*_*i*_) has the following conditional independence relation:

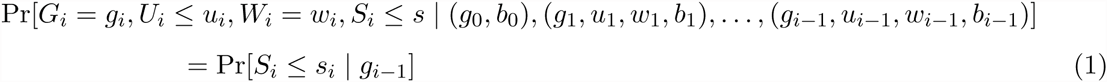

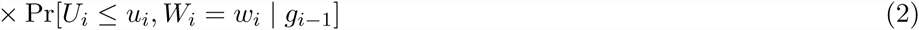

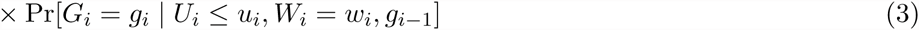

Given a chain of local genealogies, pruning locations and recombination breakpoints, the joint transition probability to a new genealogy, pruning location and locus length can be expressed as the product of the locus length probability conditioned on the current genealogy (Expression 1, above), the pruning location probability conditioned on the current genealogy (Expression 2, above) and, the transition probability of the new genealogy conditioned on the current genealogy and pruning location (Expression 3, above).

### 2.1 Complete data transition densities

Consider the chain of local genealogies **g** = (*g*_0_, *g*_1_,…,*g*_*m-*1_) with recombination breakpoints at **b** = (0, *b*_1_,…,*b*_*m-*1_). According to the SMC′ process, the first local genealogy *g*_0_ follows the standard coalescent density:

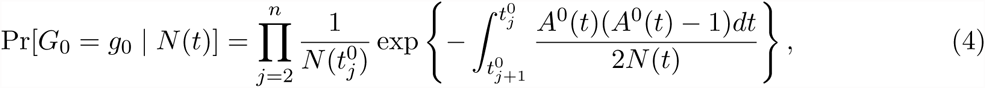

where 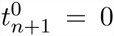 and 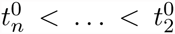are the set of coalescent times in local genealogy *g*_0_. The piece-wise constant function *A^i^*(*t*) denotes the number of ancestral lineages present at time *t* in genealogy *g*_*i*_, that is

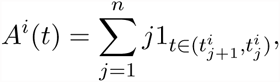

with 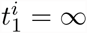.

Given a current local genealogy *g*_*i-*1_, the distribution of the length *S*_*i*_ = *B*_*i*_–*b*_*i-*1_ of the current locus depends on the current state of the SMC′ chain through the local genealogy’s total tree length *l*_*i*−1_ (the sum of all branch lengths in *g*_*i-*1_) and the recombination rate per site per generation *ρ*.

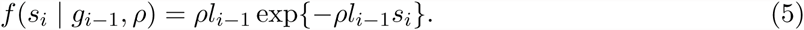

At recombination breakpoint *b*_*i*_, a new local genealogy *g*_*i*_ is generated (Figure 1). This new local genealogy *g*_*i*_ depends on the previous local genealogy *g*_*i-*1_ and the population size trajectory *N*(*t*). To generate *g*_*i*_ we first randomly choose a pruning location *p*_*i*_ (consisting of a pruning time *u*_*i*_ and a lineage *w*_*i*_) uniformly along *g*_*i-*1_. At pruning location *p*_*i*_, we add a new lineage 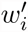 and coalesce it further in the past at time 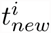 with some lineage, *c*_*i*_ (Figure 2). We then delete the *w*_*i*_ lineage’s segment from *u*_*i*_ to 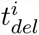 (the coalescent time of lineage *w*_*i*_). The transition density to a new genealogy at recombination breakpoint *b*_*i*_ is then

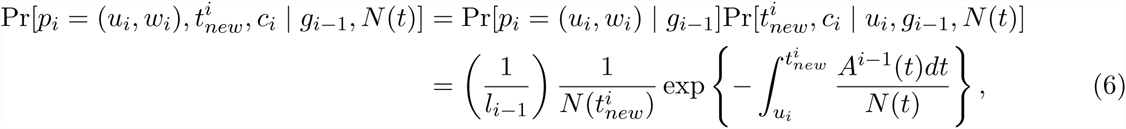

where l_*i-*1_ denotes the total tree length of *g*_*i-*1_.

This generative process of local genealogies can result in the two types of transitions depicted in Figure 2. A *visible transition* results in a genealogy *g*_*i*_ which is different from *g*_*i-*1_ (Figure 2A), while an *invisible transition* makes *g*_*i*_ identical to *g*_*i-*1_ (Figure 2B).

**Figure 2:**
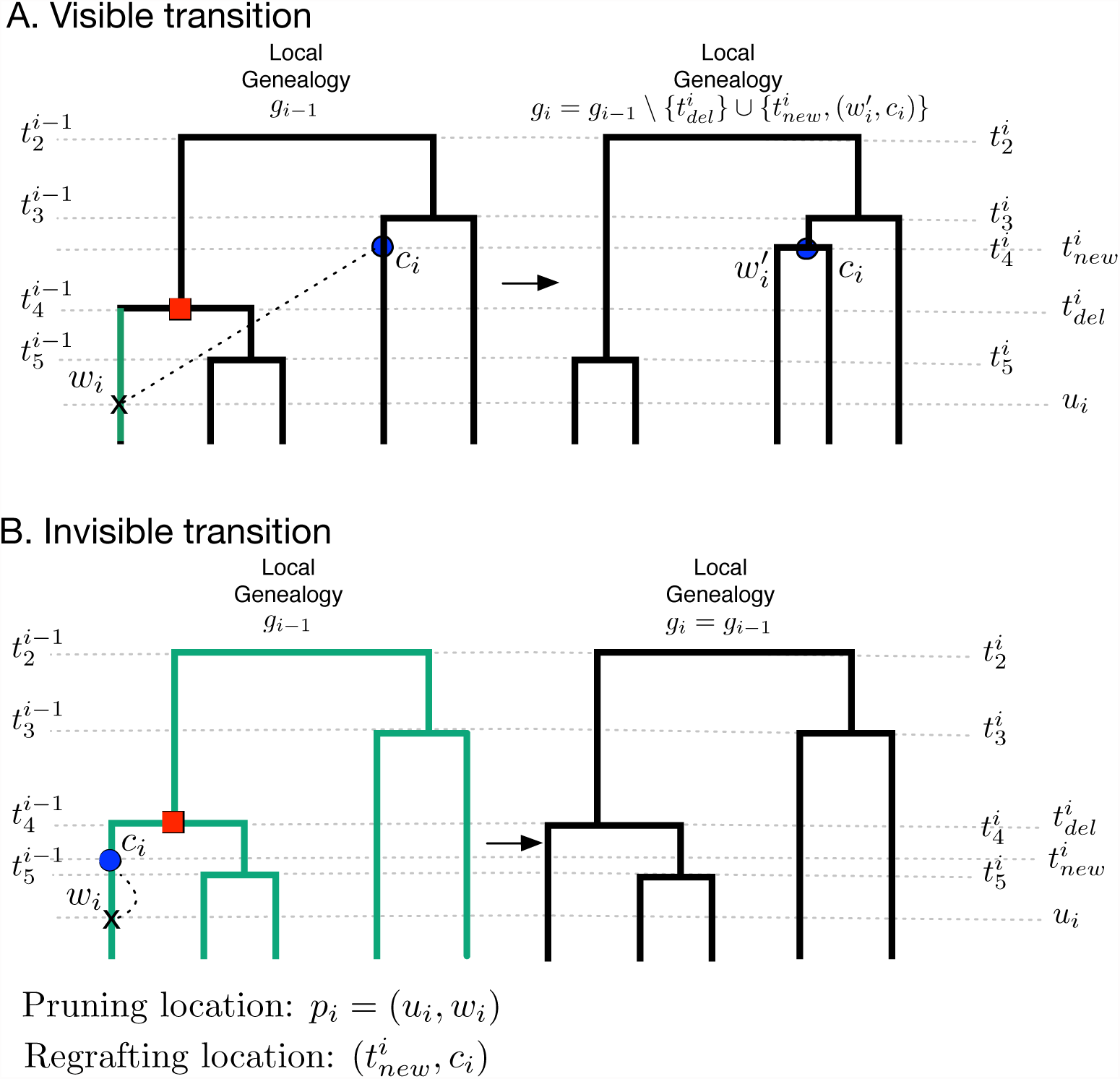
Schematic representation of SMC′ transitions given a recombination break-point at location *b*_*i*_ (indicated as an arrow in each panel). **A: Visible transition.** We uniformly sample the pruning location *p*_*i*_ from *g*_*i-*1_ at time *u*_*i*_ along some branch *w*_*i*_, we add a new branch 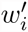 at *u*_*i*_ and re-graft it (dashed black line). The new branch 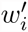 coalesces with some branch *c*_*i*_ at time 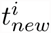. We then delete branch *w*_*i*_ and the coalescent time 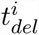 to generate genealogy *g*_*i*_. Any pruning time along the branch *w*_*i*_ (shown in green) would have produced the same visible transition from *g*_*i-*1_ to *g*_*i*_. **B: Invisible transition.** We uniformly sample the pruning location *p*_*i*_ = (*u*_*i*_, *w*_*i*_), add a new branch 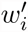 at *u*_*i*_ and re-graft it. The new branch 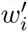 coalesces with itself (dashed black line); that is, *C*_*i*_ = *w*_*i*_, and then the segment 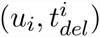 of *w*_*i*_ is deleted. If *C*_*i*_ = *w*_*i*_, any pruning location along the green branches would have produced the same invisible transition.

An invisible transition *g*_*i*_ = *g*_*i-*1_, occurs when *c*_*i*_ = *w*_*i*_. Given the pruning location *p*_*i*_ = (*u*_*i*_, *w*_*i*_), a transition to an invisible event occurs when 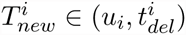 and *C*_*i*_, the random variable indicating the lineage that coalesces with lineage 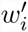, takes the value *w*_*i*_. The probability of an invisible transition is given by

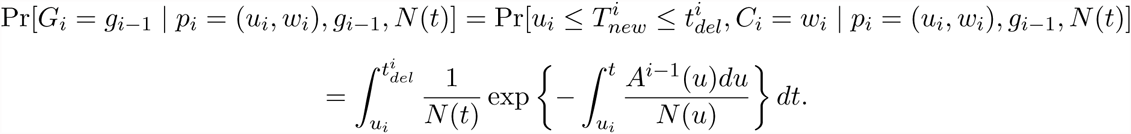

Thus, the joint transition probability to an invisible event with pruning location (*u*_*i*_, *w*_*i*_), given *g*_*i-*1_is:

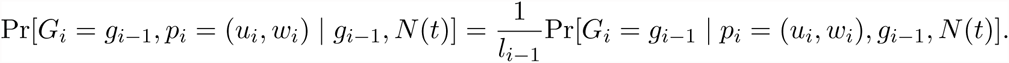

### 4.2 Transition densities averaged over unknown pruning locations

Even though we will assume that local genealogies are known, in order to anticipate later applications to sequence data we do not wish to make the same assumption about pruning locations. Thus, we average over pruning locations to obtain marginal transition densities between genealogies, for both visible and invisible transitions.

To compute the marginal visible transition density to a new genealogy 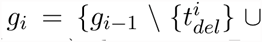 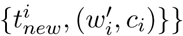, we need to average over all possible pruning locations *p*_*i*_ = (*u*_*i*_, *w*_*i*_) along *g*_*i-*1_. By comparing the two genealogies *g*_*i-*1_ and *g*_*i*_ in Figure 2A, we know that *p*_*i*_ corresponds to the lineage *w*_*i*_ some time along 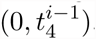, or equivalently, along 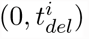. In general, comparison of *g*_*i-*1_ and *g*_*i*_may not provide complete information to identify the lineage that was pruned. When the children of the node corresponding to *t*_*del*_ and the children of the node corresponding to *t*_*new*_ are the same, pruning different branches can lead to the same transition. We enumerate all cases of incomplete information for visible transitions in Supporting Information Figure S1.

We introduce a function *I*^*i-*1^(*t*), equal to the number of possible lineages at time *t* where the pruning location along *g*_*i-*1_ would produce a visible transition to *g*_*i*_. *I*^*i-*1^(*t*) is a piece-wise constant function that takes the values in {0, 1, 2} depending on whether the pruning location *p*_*i*_ can happen in 0, 1 or 2 branches at time t. In the example in Figure 2A,

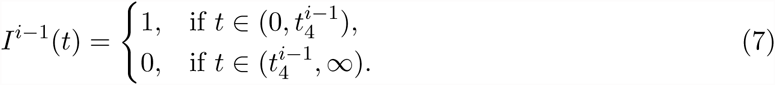

For a general *I*^*i-*1^(*t*) piece-wise constant function that indicates the number of possible pruning branches at time *t*, the marginal visible transition density to a new genealogy is

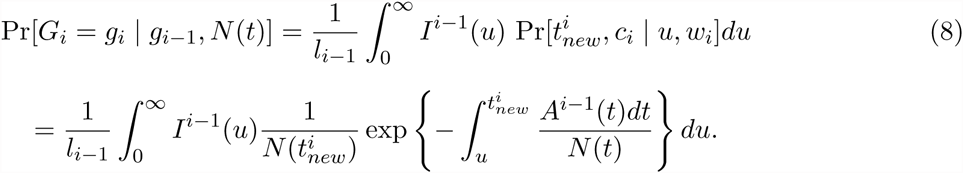

Turning now to the computation of marginal transition probabilities for invisible events, we need to average over all possible pruning locations *p*_*i*_. Consider the example in Figure 2B and choosing a pruning time (*u*_*i*_) along *g*_*i-*1_. In order to have an invisible transition, the coalescing branch *C*_*i*_ must be the same pruning branch *W*_*i*_. In Figure 2B the new coalescent time 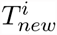 can happen along five lineages in the interval 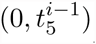, three lineages in the interval 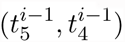, and two lineages in the interval 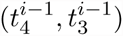. To generalize this calculation, we introduce the quantity 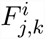 with (*n* + 1) ≥ *j* ≥ *k* ≥ 2 which denotes the number of lineages in *g*_*i*_ that are *free* (do not coalesce), in the time segment 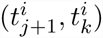. The time interval 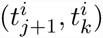 includes the interval of pruning 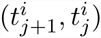 up to the interval of self-coalescence 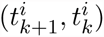. Thus, if the pruning time happens at time*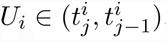*, an invisible transition with new coalescent time*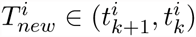* can happen along 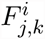 free lineages.

In Figure 2B, *u*_*i*_ happened in the time interval 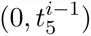. If the new coalescent time *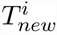*happens in the interval 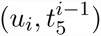 along the same (unknown) pruning branch, then this invisible transition has probability

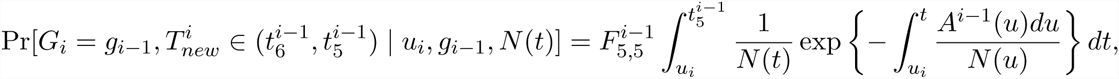

with *F*_5,5_ = 5.

Now consider the same example of Figure 2B but with an unknown pruning time *u*_*i*_. The joint event where recombination occurs at pruning time 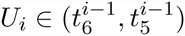 and coalescent time *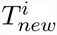* occurs in the interval 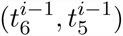 and this results in an invisible transition has probability:

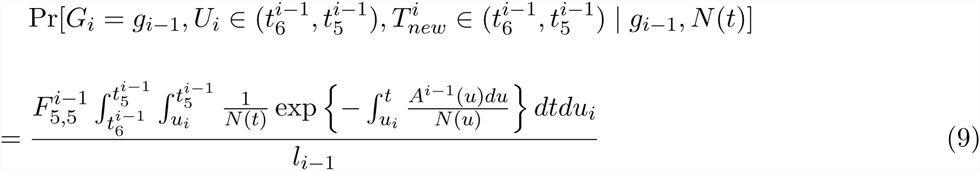

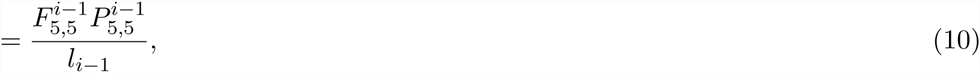

where 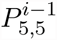 denotes the double integral expression in Equation 9 for ease of notation.

An invisible transition would also result if 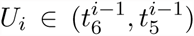 and *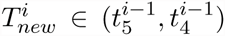* along the same (unknown) pruning branch; according to Figure 2B, this can happen along three lineages, so 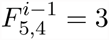 and this event has probability:

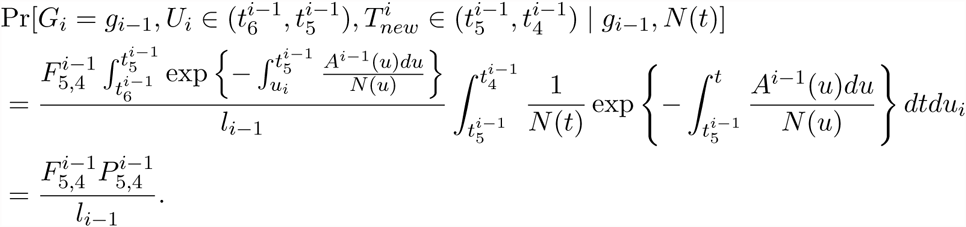

If we continue considering the cases where 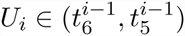 and *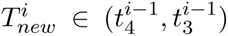* or 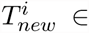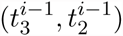 we have 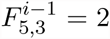 and 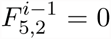. Then, the joint probability of an invisible event and 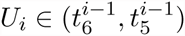 is

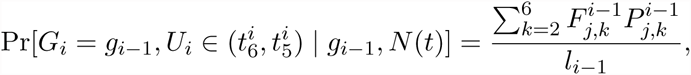

For the cases when 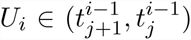 and the new coalescent time *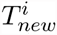* falls in another coalescent interval 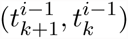, we need to compute the following:

- The joint probability of 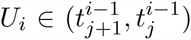 and no coalescence in the interval 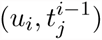:

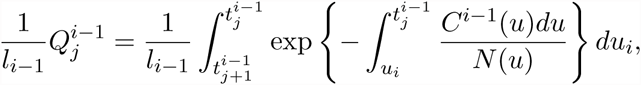
- The probability of no coalescence in any of the intermediate coalescent intervals 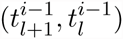:

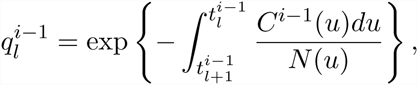

and
- The probability of coalescing at 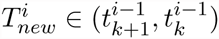

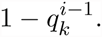

Then,

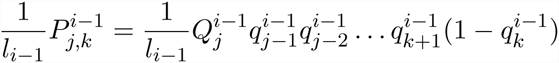

represents the probability that the pruning location is *w*_*i*_ at time 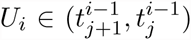 and the new lineage 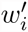 coalesces at time *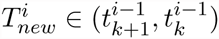* with lineage *c*_*i*_ = *w*_*i*_. Overall, the marginal transition probability to an invisible event is:

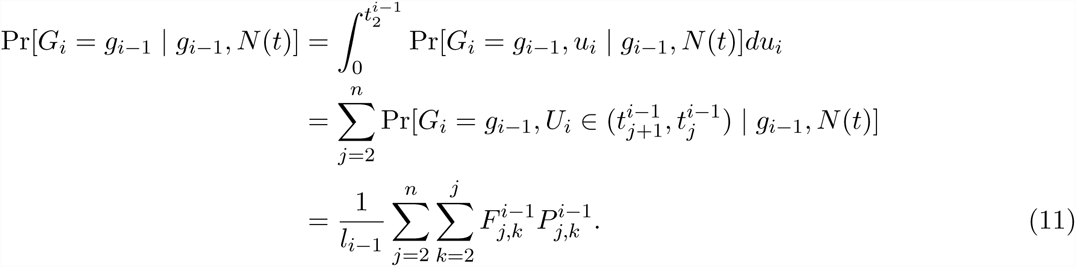

### 2.3 The likelihood of the embedded SMC′ chain

Instead of having a complete realization of the embedded SMC′ chain of *m* local genealogies *g*_0_,…,*g*_*m-*1_ and pruning locations *p*_1_,…,*p*_*m-*1_ at recombination breakpoints *b*_1_,…,*b*_*m-*1_, we assume that our data (unless otherwise noted) consist only of *m* local genealogies at recombination breakpoints from a chromosomal segment of length *L*. Note that our observed data are not sequence data. More specifically, our observed data are

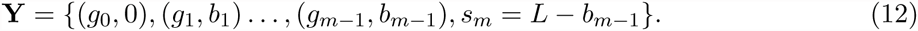

Then, the observed data likelihood is

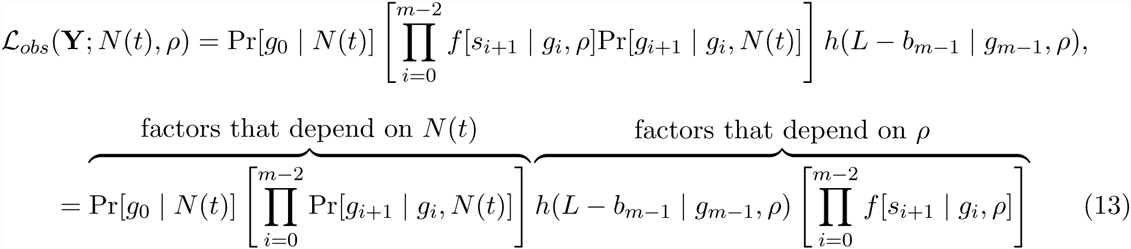

where *h*(*L* − *b*_*m-*1_ | *g*_*m-*1_, *ρ*) is the survival function in state *g*_*m-*1_. Equation 13 is factored into terms that depend on *N*(*t*) alone and ones that depend on *ρ* alone. The terms that depend on *ρ*, given by Equation 5, depend on the data only through total tree lengths *l*_0_,…, *l*_*m*−1_ and locus lengths *s*_1_,…, *s*_*m*-1_, *L* − *b*_*m*−1_. By the factorization theorem for sufficient statistics, local tree lengths *l*_0_,…, *l*_*m*−1_ and locus lengths *s*_1_,…, *s*_*m*−1_, *L* − *b*_*m*-1_ are sufficient for inferring *ρ*.

## Methods: Inference

Current coalescent-based methods that infer a population size trajectory *N*(*t*) from whole-genome data assume *N*(*t*) is a piece-wise constant function with change points *x*_1_ = 0 < *x*_2_ <… < *x*_*d*_ (Li and Durbin 2011; Sheehan et al. 2013; Rasmussen et al. 2014; Schiffels and Durbin 2014). That is

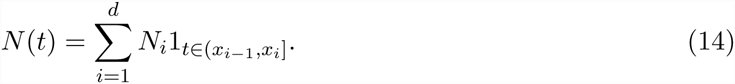

Equation 14 presents two challenges. The first challenge lies in the specification of the change points. The narrower an interval is, the higher the probability that we do not observe coalescent times in that interval. The fewer observed coalescent times in an interval, the greater the uncertainty of the estimate 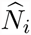 (if the estimate even exists). The second challenge lies in the specification of the time window (0, *x*_*d*_): if *x*_*d*_ is set too far in the past, we might not have enough data to accurately estimate *N*(*t*) for *x*_*d*_≤*t* < ∞.

In order to solve the first challenge, Rasmussen et al. (2014) and Li and Durbin (2011) distribute the *d* change points evenly on a logarithmic scale:

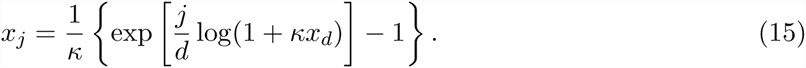

where κ is specified by the user. Schiffels and Durbin (2014) propose discretizing time according to the quantiles of the exponential distribution.

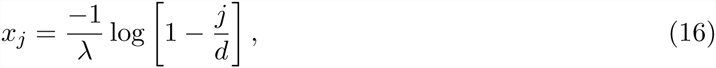

where λ is the rate of an exponential distribution. Schiffels and Durbin (2014) model the time to the most recent coalescent event and set 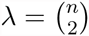. However, Equation 16 is not directly applicable here because we use all coalescent events for inference.

In the following sections, we first present our Bayesian nonparametric method, then develop a maximum likelihood method under a piece-wise constant trajectory so we can directly compare an EM-based method (Li and Durbin 2011; Sheehan et al. 2013) to our Bayesian nonparametric method. The R code for all simulation studies and real data analysis conducted in this paper are publicly available at http://ramachandran-data.brown.edu/datarepo/.

### 3.1 Gaussian-Process-based Bayesian Nonparametric Estimation of *N*(*t*)

For our Bayesian methodology, we assume the following log-Gaussian Process prior on the population size trajectory, *N*(*t*):

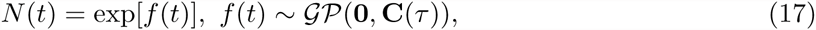

where **𝒢 𝒫**(**0**, C(τ) denotes a Gaussian process with mean function **0** and inverse covariance function **C**^-1^(τ) = τ **C**^-1^ with precision parameter τ. For computational convenience, we use Brownian motion as our prior for *f*(*t*) since its inverse covariance matrix is sparse. We place a Gamma prior on the precision parameter τ,

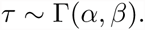

Assuming that recombination rate *ρ* is known, the posterior distribution of model parameters (Figure 3) is then

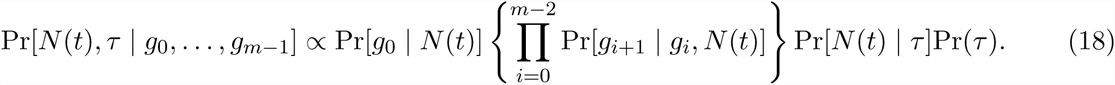

The first two factors on the right side of Equation 18, detailed in Equations 8 and 11 involve integration over *N*(*t*), an infinite dimensional random function (Equation 17). We approximate the integral

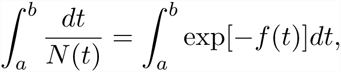

by the Riemann sum over a partition of the integration interval. That is,

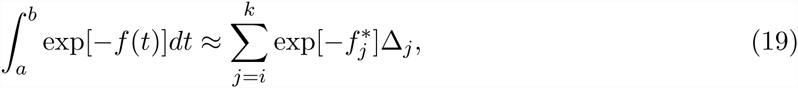

for *x*_*i*_ < *a* < *x*_*i*+1_ <… < *x*_*k-*1_ < *b* < *x*_*k*_, Δ*_i_* = *x*_*i*+1_ − *a*, Δ*_k_* = *b* − *x*_*k*-1_ and Δ*_j_* = *x*_*j*+1_ − *x*_*j*_ for *i* < *j* < *k*. 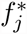 is a representative value of *f*(*t*) in the interval (*x*_*j*_, *x*_*j*+1)_; in our implementation, we set 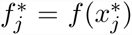 with 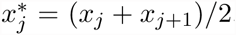. This way, we discretize our time window in *d* evenly spaced segments *x*_1_ = 0 < *x*_2_ <… < *x*_*d*_, with 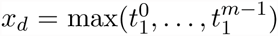, the maximum time to the most common ancestor observed in the sequence of local genealogies, and approximate *N*(*t*) by a piece-wise linear function evaluated at 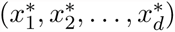.

**Figure 3:**
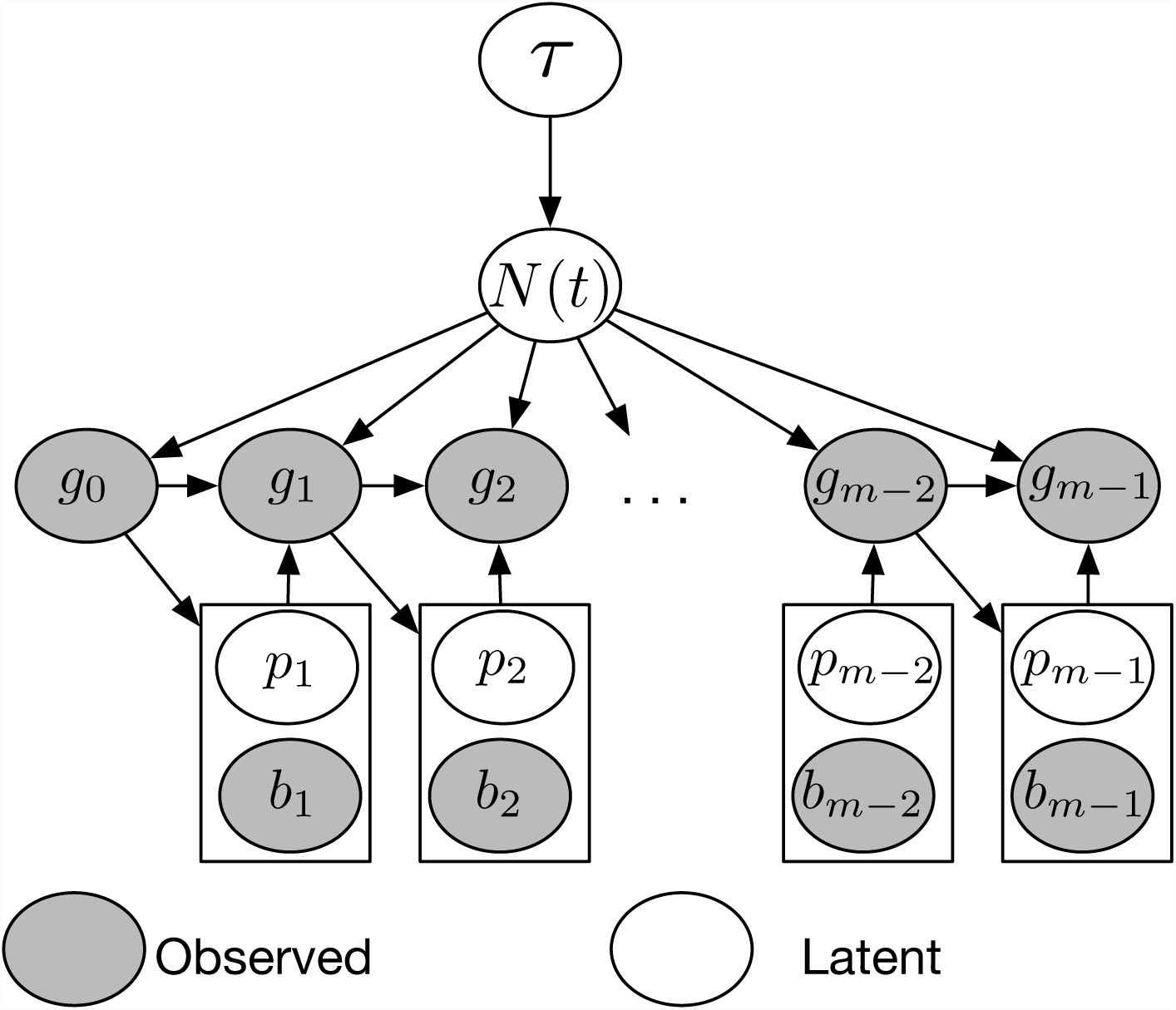
Structure of our Bayesian model for inferring population size trajectories from a realization of the SMC′ process at recombination breakpoints. Hyperparameter τ controls the smoothness of the log-Gaussian process prior on *N*(*t*). Local genealogies depend on *N*(*t*) and form a Markov chain of degree one. Given the current local genealogy *g*_*i-*1_, we sample the location of the new recombination breakpoint *b*_*i*_ and a pruning location *p*_*i*_ on genealogy *g*_*i-*1_. The new genealogy *g*_*i*_ depends on *N*(*t*), *p*_*i*_ and *g*_*i-*1_.

We condition on the set of *m* local genealogies *g*_0_,…,*g*_*m-*1_ to generate posterior samples for the vector 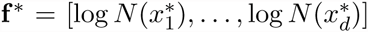 and τ and use these posterior samples to infer *N*(*t*) at *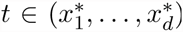*, where 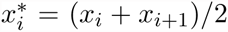. Updating *N*(*t*) and τ separately is not recommended because of their strong dependency (Lan et al. 2015). Therefore, we update (*N*(*t*), τ) jointly in an MCMC sampling algorithm using Split Hamiltonian Monte Carlo (Shahbaba et al. 2014; Lan et al. 2015). Split Hamiltonian Monte Carlo relies on our ability to calculate the log-likelihood of the observed data and the gradient vector of the log-likelihood (i.e., the score function). The log-likelihood of the observed data is approximated via sums of the form in Equation 19. We approximate the score function ∇ℒ_*obs*_(**Y**; **f\***) with respect to **f\*** by applying Fisher’s identity:

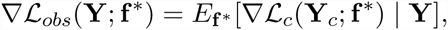

where, at each iteration in the MCMC, expectation is calculated using the current value of **f\***. We show the details of this calculation in the Appendix.

Alternatively, one can update *N*(*t*) in the MCMC algorithm using Elliptical Slice Sampler (Murray et al. 2010) with a fixed value of τ (perhaps estimated from previous studies or from a preliminary run from the Split Hamiltonian Monte Carlo algorithm). The advantage of using Elliptical Slice Sampler over the Split Hamiltonian Monte Carlo is purely computational since Elliptical Slice Sampler does not require calculation of the score function.

### 3.2 Maximum-likelihood estimation of *N*(*t*) with measures of uncertainty

We assume that the population size trajectory *N*(*t*) is defined as in Equation 14. The standard coalescent density (Equation 4) and the transition densities defined in Equations 11 and 8 are tractable, so calculation of the likelihood (Equation 13) is tractable. However maximization of the likelihood function cannot be performed analytically because pruning locations are missing. We rely on the Expectation-Maximization (EM) algorithm (Dempster et al. 1977) to find the maximum likelihood estimator of **N** = (*N*_1_,…,*N*_*d*_). The complete data **Y***_c_* for inferring *N*(*t*) are then the set of local genealogies *g*_0_,…,*g*_*m-*1_ and the set of pruning locations *p*_1_…, *p*_*m-*1_. For the invisible transitions, we also need to know the new coalescent times 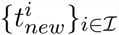, where ℐ ⊂ {1, 2,…,*m* − 1} denotes the set of indices of invisible transitions (transition *i* is an invisible transition if *g*_*i*_ = *g*_*i-*1_).

The complete data log-likelihood is then

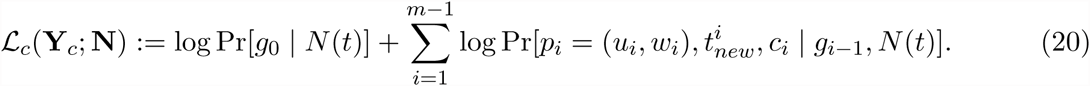

The EM algorithm starts by initializing the population size trajectory to a piece-wise constant function with change points *x*_1_,…, *x*_*d*_ with arbitrarily chosen vector **N**^0^. At the *k*th iteration of the algorithm we set

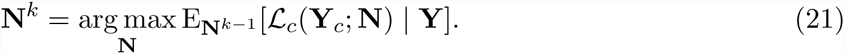

The conditional expectation in Equation 21 is conditional on the observed data **Y** defined in Equation 12. Let 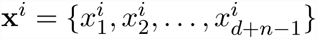 be the ordered set of time points corresponding to the change points *x*_1_,…,*x*_*d*_ and the coalescent time points **t^*i*^** of local genealogy *i*. If the transition from *g*_*i*_ to *g*_*i*+1_ is visible, we replace the jth time point 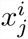 by 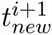, where *j* corresponds to the index such that 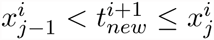. For ease of notation, we will denote the number of time intervals |**x***^i^*| by *D* = *d* + *n* − 2. Let

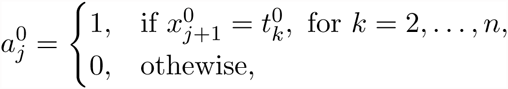

be an indicator function that takes the value of 1 when the *j*th interval contains a coalescent time of the first genealogy *g*_0_. Then, the log density of the first genealogy is:

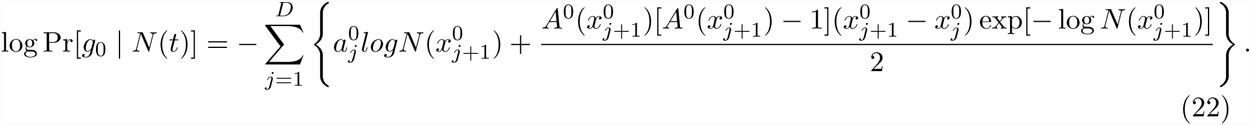

Let

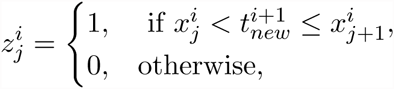

be an indicator function that takes the value of 1 when the new coalescent time of genealogy *i* happens in the corresponding time interval 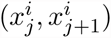, and let the adjusted interval length be

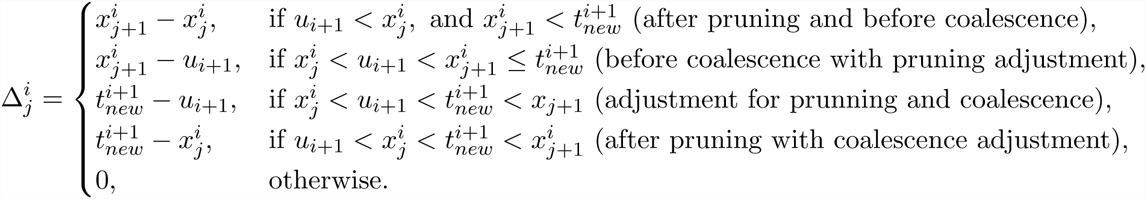

Then, the augmented transition density can be expressed as:

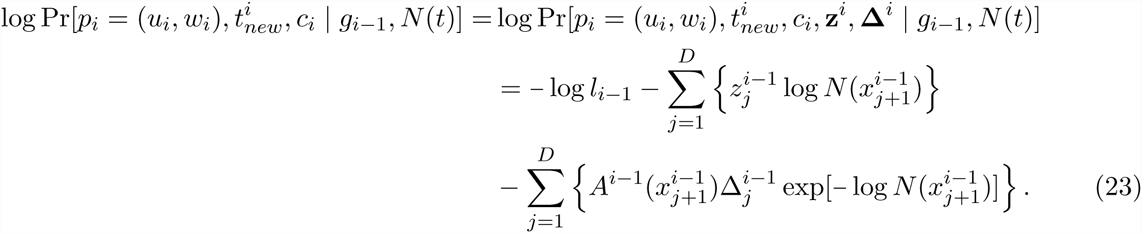

where **z***^i^* and **Δ** *^i^* are the vectors with 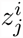 and 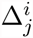 elements. For the EM algorithm we need to compute the conditional expected vectors 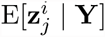 and 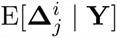. The details of these calculation are in the Appendix.

We use the Fisher information matrix to compute approximate standard errors of 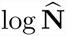 and use these standard errors together with asymptotic normality of maximum likelihood estimators to produce confidence intervals for log population size piece-wise trajectories. We compute the observed Fisher information matrix following Louis (1982):

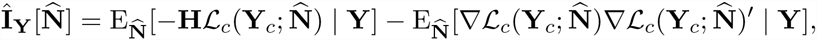

where 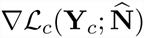 is the gradient and 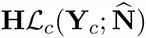 is the Hessian of the complete-data log-likelihood with respect to log **N**. This requires the calculation of conditional cross-product means and conditional second moments described in the Appendix.

## Results

We simulated 1000 local genealogies of 2, 20 and 100 individuals from each of the three different demographic models described in Table 2 using `MaCS` (Chen et al. 2009); see Supporting Information for details of these simulations. We assumed that all individuals were sampled at time *t* = 0 under a demographic model in Table 2.

**Table 2:** Simulated demographic scenarios. The argument *t* denotes time measured in units of *N*_0_ generations.

| Demographic model | $N(t)$ |
| --- | --- |
| Constant Population size: | $N(t) = 1$ |
| Exponential growth followed by constant size: | $N(t) = \begin{cases} 1, & \text{for } t \in (0, 0.1), \\ \exp[-10(t - 0.1)], & \text{for } t \in (0.1, \infty). \end{cases}$ |
| Population bottleneck: | $N(t) = \begin{cases} 1, & \text{for } t \in (0, 0.3), \\ 0.1, & \text{for } t \in (0.3, 0.5), \\ 1, & \text{for } t \in (0.5, \infty). \end{cases}$ |

We compared the point estimates with the truth for each demographic model using the sum of relative errors (SRE):

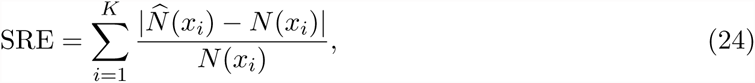

where *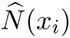* is the estimated population size trajectory at time *x*_*i*_. We compute SRE at equally space time points *x*_1_,…,*x*_*K*_. Second, we compute the mean relative width (MRW) as follows:

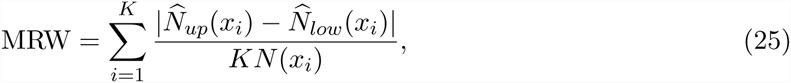

where 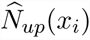 corresponds to the 97.5% upper limit and 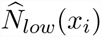 corresponds to the 2.5% lower limit of *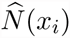*. For EM estimates, 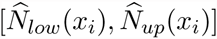 corresponds to the 95% confidence interval estimated using the observed Fisher information; for Bayesian GP estimates, 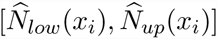 corresponds to the 95% Bayesian credible interval (BCI) of *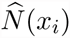*. To measure how well these intervals cover the truth, we compute the envelope measure (ENV) in the following way:

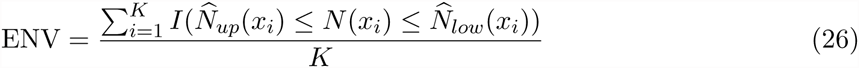

We compute SRE, MRW and ENV for *K* = 150 at equally spaced time points.

For our Bayesian GP estimates, we estimate *N*(*x*_*i*_) at *d* = 100 time points, unless stated otherwise. The parameters of the Gamma prior on the GP precision parameter τ were set to *α* = *β* = 0.001, reflecting our lack of prior information about the smoothness of the population size trajectory.

For our EM estimates, we used different discretizations based on Equation 15 and varying the number of change points *d* and κ over the fixed interval (0, *x*_*d*_) with *x*_*d*_ set to be the maximum observed coalescent time. For the cases where we only consider one genealogy (*m* = 1), the EM approach becomes standard maximum likelihood estimation. We summarize our posterior inference and compare our Bayesian GP method to the EM method. The population size trajectory is logtransformed for ease of visualization and for direct comparison with other methods (Minin et al. 2008; Palacios and Minin 2013).

**Table 3:**
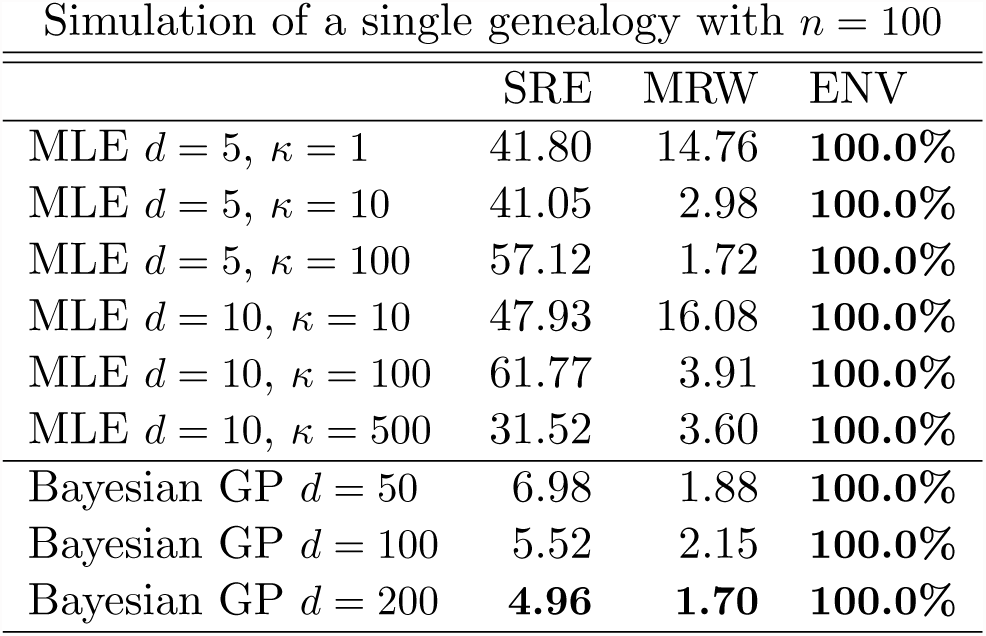
Summary statistics for simulation results depicted in Figure 4. SRE is the sum of relative errors (Equation 24), MRW is the mean relative width of the 95% BCI (Equation 25), and ENV is the envelope measure (Equation 26). Values in bold indicate best performance.

| Simulation of a single genealogy with $n = 100$ | | | |
| --- | --- | --- | --- |
|  | SRE | MRW | ENV |
| MLE $d = 5, \kappa = 1$ | 41.80 | 14.76 | **100.0%** |
| MLE $d = 5, \kappa = 10$ | 41.05 | 2.98 | **100.0%** |
| MLE $d = 5, \kappa = 100$ | 57.12 | 1.72 | **100.0%** |
| MLE $d = 10, \kappa = 10$ | 47.93 | 16.08 | **100.0%** |
| MLE $d = 10, \kappa = 100$ | 61.77 | 3.91 | **100.0%** |
| MLE $d = 10, \kappa = 500$ | 31.52 | 3.60 | **100.0%** |
| Bayesian GP $d = 50$ | 6.98 | 1.88 | **100.0%** |
| Bayesian GP $d = 100$ | 5.52 | 2.15 | **100.0%** |
| Bayesian GP $d = 200$ | **4.96** | **1.70** | **100.0%** |

**Figure 4:**
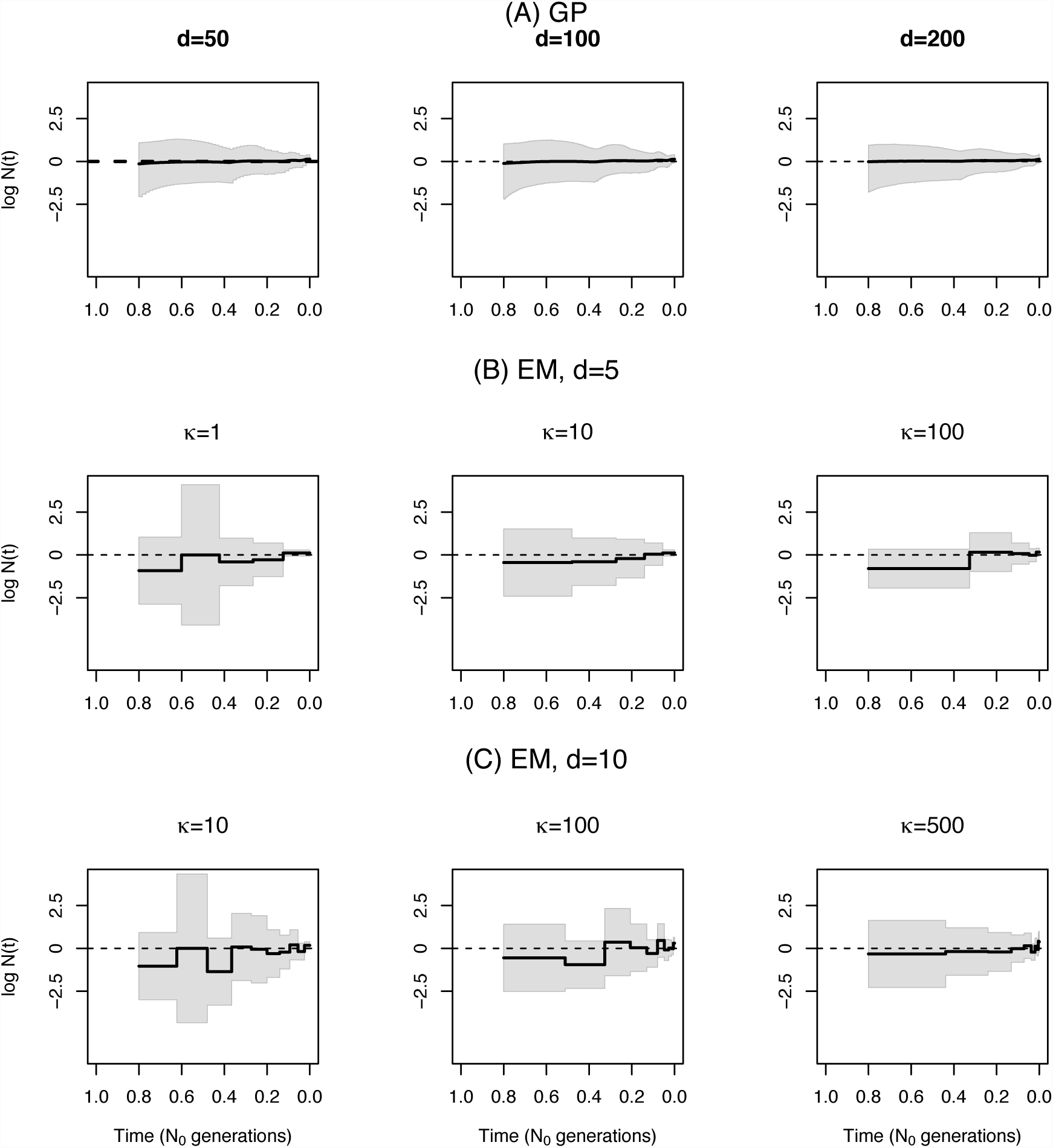
Sensitivity to parameter discretization. Comparison of population size trajectories estimated from one simulated genealogy (*m* = 1) of 100 individuals with a constant population size. We show true trajectories as dashed lines. (A) Bayesian GP estimates at *d* = 50, 100 and 200 equally spaced time points. (B) EM estimates of a piece-wise constant trajectory with *d* = 5 change points and *κ* = 1, 10 and 100 (Equation 15). (C) EM estimates of a piece-wise constant trajectory with *d* = 10 change points and *κ* = 10, 100 and 500 (Equation 15). Point estimates are shown as solid black lines. 95% credible intervals and 95% confidence intervals are shown by gray shaded areas.

### 4.1 Sensitivity of EM estimates of *N* (*t*) to discretization

In Figure 4, we show our Bayesian GP and EM estimates of a constant population size trajectory from a single genealogy of 100 individuals with different discretizations. We find that our Bayesian GP point estimates depicted in Figure 4A recover the truth (dashed line) almost perfectly with less uncertainty than the EM (Figure 4B-C). Comparing our Bayesian GP estimates with different discretizations: 50, 100 and 200 equally spaced time points (Figure 4A), we find that increasing the number of time points improves inference (Table 4) but that the differences between estimates among the three discretizations are marginal (Figure 4A). In contrast, we show that different grid definitions alter the EM estimates (Figure 4B). It is not clear how to define a good strategy for the definition of the grid for the EM method, even for the simple model of constant population size. For example, increasing *κ* from 100 to 500 with 5 change points (Figure 4B), does not improve estimation. Increasing the number of change points does not necessarily improve the estimates either; for example, increasing the the number of change points from 5 to 10 for *κ* = 10 (Figures 4B-C). EM grid sensitivity is persistent even when the number of genealogies increases; Figure S2 in Supplementary Information shows that the best definition of change points when our data consist of 1000 local genealogies of 100 individuals has 10 change points evenly distributed.

**Table 4:** Summary of simulation results depicted in Figures 5. SRE is the sum of relative errors calculated as in (24), MRW is the mean relative width of the 95% BCI as defined in (25), and ENV is the envelope measure calculated as in (26). Values in bold indicate best performance for each demographic model and sample size.

| A. Simulations with $n = 2$ | | | | | | | | | |
| --- | --- | --- | --- | --- | --- | --- | --- | --- | --- |
|  | SRE |  |  | MRW |  |  | ENV |  |  |
|  | m=100 | m=500 | m=1000 | m=100 | m=500 | m=1000 | m=100 | m=500 | m=1000 |
| Const. EM | 39.80 | 41.78 | 38.60 | 0.98 | **0.26** | **0.08** | 31.3% | 28.0% | 19.3% |
| Const. GP | **30.60** | **4.25** | **3.04** | **0.49** | 0.33 | 0.22 | **100.0%** | **100.0%** | **100.0%** |
| Exp. EM | 64.68 | **25.70** | 33.70 | **0.91** | **0.16** | **0.12** | 42.0% | 26.0% | 6.6% |
| Exp. GP | **28.38** | 32.70 | **26.76** | 2.04 | 0.45 | 0.33 | **100.0%** | **56.0%** | **50.6%** |
| Bottle. EM | 48.48 | 46.51 | **127.70** | **0.43** | **0.45** | **1.37** | 40.6% | 30.0% | 34.0% |
| Bottle. GP | **33.76** | **45.14** | 223.58 | 3.44 | 6.84 | 17.13 | **98.0%** | **94.6%** | **94.6%** |

| B. Simulation with $n = 20$ | | | | | | | | | |
| --- | --- | --- | --- | --- | --- | --- | --- | --- | --- |
|  | SRE |  |  | MRW |  |  | ENV |  |  |
|  | m=1 | m=100 | m=1000 | m=1 | m=100 | m=1000 | m=1 | m=100 | m=1000 |
| Const. EM | 60.87 | 121.30 | 25.60 | 2.28 | 2.16 | 0.23 | **100.0%** | 37.7% | 39.3% |
| Const. GP | **31.74** | **3.94** | **13.22** | **1.06** | **0.70** | 0.36 | **100.0%** | **100.0%** | **100.0%** |
| Exp. EM | 40.97 | 40.66 | **40.22** | **3.11** | **0.37** | **0.19** | **100.0%** | 38.6% | 19.3% |
| Exp. GP | **25.35** | **27.03** | 65.61 | 3.53 | 1.56 | 0.42 | **100.0%** | **100.0%** | **39.3%** |
| Bottle. EM | 147.93 | 78.40 | 78.20 | 6.98 | **0.81** | 68.4 | 66.0% | 78.6% | 49.33% |
| Bottle. GP | **68.93** | **78.2** | **50.92** | **2.74** | 2.47 | **1.47** | **92.0%** | **79.3%** | **78.6%** |

| C. Simulation with $n = 100$ | | | | | | | | | |
| --- | --- | --- | --- | --- | --- | --- | --- | --- | --- |
|  | SRE |  |  | MRW |  |  | ENV |  |  |
|  | m=1 | m=100 | m=1000 | m=1 | m=100 | m=1000 | m=1 | m=100 | m=1000 |
| Const. EM | 41.05 | 220.85 | 43.41 | 2.98 | 4.93 | 0.99 | **100.0%** | 35.3% | 48.0% |
| Const. GP | **5.52** | **34.78** | **12.17** | **2.15** | **1.49** | **0.47** | **100.0%** | **100.0%** | **89.3%** |
| Exp. EM | **76.86** | 40.22 | 27.63 | **3.23** | **0.81** | **0.13** | 87.3% | 42.0% | 14.0% |
| Exp. GP | 114.53 | **25.82** | **26.42** | 3.57 | 1.55 | 0.83 | **100.0%** | **100.0%** | **100.0%** |
| Bottle. EM | 194.77 | 59.54 | 127.68 | **3.95** | **1.08** | **0.85** | 84.0% | 51.3% | 45.3% |
| Bottle. GP | **90.27** | **44.14** | **42.68** | 6.98 | 2.62 | 1.74 | **100.0%** | **94.7%** | **96.0%** |

### 4.2 Comparing Methods of Estimating *N* (*t*)

Figure 5 shows the estimated population size trajectories when the number of samples is 2 for the three different demographic scenarios and varying the number of local genealogies (100, 500 and 1000 local genealogies). For constant and exponential growth, our EM method assumes a piece-wise constant trajectory of 10 change points (*d* = 10) and *κ* = 1 using Equation 15 (similar to Li and Durbin (2011) and Rasmussen et al. (2014)). For the bottleneck scenario, some of the intervals did not have coalescent events; hence, for this case we assumed a piece-wise constant trajectory of 5 change points (*d* = 5) and *κ* = 1 for constructing our EM estimates. We show the boxplots of the time to the most recent common ancestor (TMRCA) at the bottom of each plot in Figure 5. The distribution of the TMRCA serves as an indicator of the uncertainty expected of our estimates. Both approaches, EM and Bayesian GP show narrower confidence and credible intervals at the center of the distribution of the TMRCA, particularly during the bottleneck in Figure 5C.

**Figure 5:**
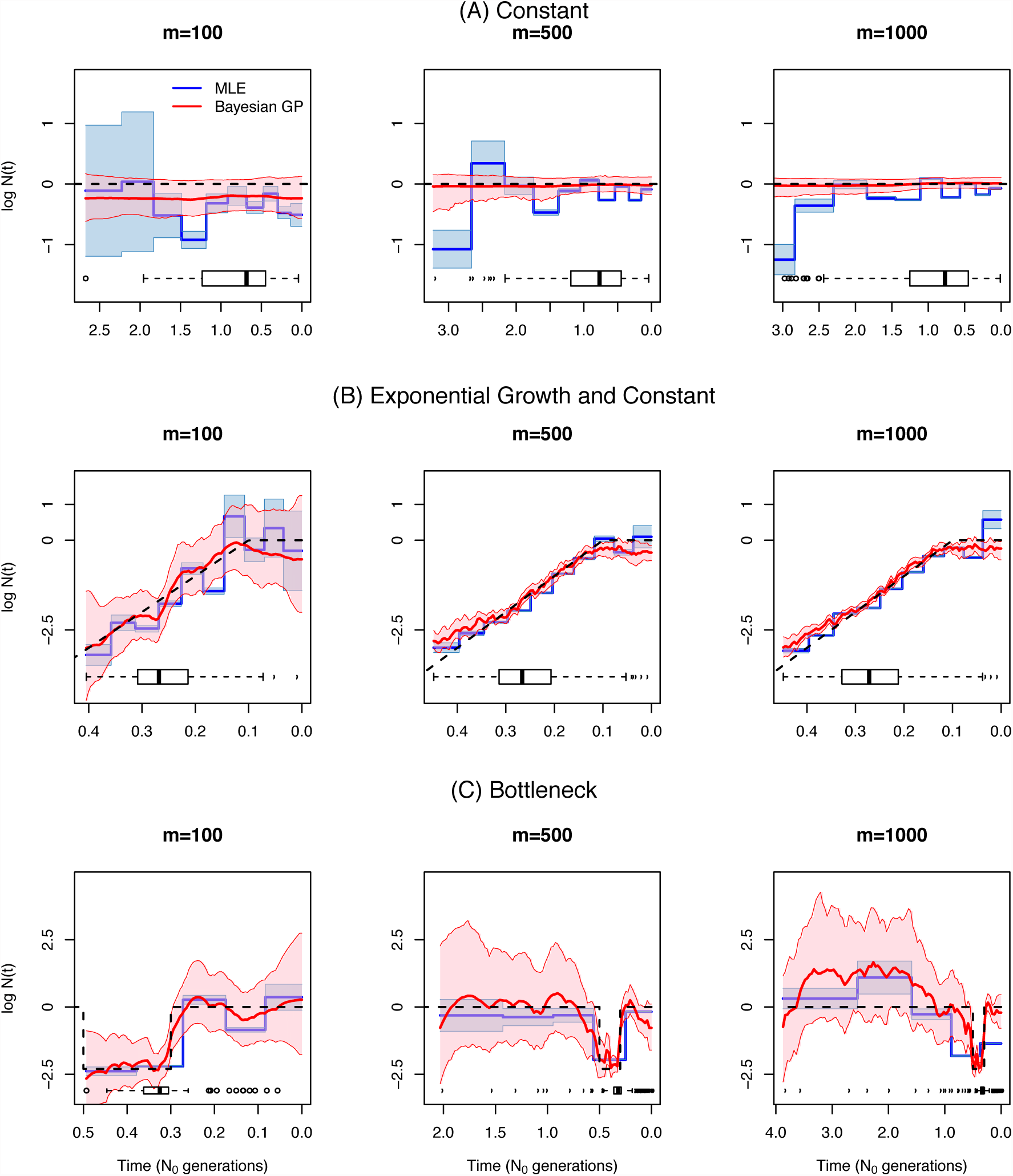
Inference of population size trajectories *N*(*t*) for a pair of individuals (*n* = 2). (A) Simulated data under constant population size, (B) exponential and constant trajectory, and a bottleneck. We show estimates from *m* = 100, *m* = 500, and *m* = 1000 local genealogies. We show the true trajectories as dashed lines, blue lines and light blue shaded areas represent EM point estimates and 95% confidence areas, and red lines and pink shaded areas represent Bayesian GP posterior medians and 95% BCIs. Boxplots of the TMRCA are shown at the bottom of each plot.

For the constant population demographic model in Figure 5A, our Bayesian GP outperforms our EM estimates considerably. This is not surprising since *a priori* log *N*(*t*) has mean 0 in our Bayesian approach (Equation 17). Moreover, EM confidence intervals only cover the truth constant population size around 30% of the time, while the GP method covers 100% of the truth (Table 4A). Despite placing a mean-0 prior on *logN*(*t*), the Bayesian GP method accurately recovers sudden changes as shown in the bottleneck example. Our Bayesian GP prior on log *N*(*t*) is Brownian motion which is not differentiable at any point; yet, our Bayesian GP recovers smooth curves (Figure 5B).

Table 4A shows the performance statistics for the estimates of *N*(*t*) in Figure 5. In general, our Bayesian GP has wider credible intervals than the EM confidence intervals but these credible intervals cover the true trajectory better than the EM confidence intervals in all the cases (MRW and ENV in Table 4). Our Bayesian GP estimates also generally have smaller sums of relative errors (SRE in Table 4). Under the bottleneck scenario, our Bayesian GP produces greater sums of relative errors than does the EM, but our Bayesian GP estimates recover the truth more accurately than the EM during the bottleneck.

Figures 6 and 7 show our estimates when *n* = 20 and *n* = 100. The performance statistics of the estimates displayed are shown in Table 4(B) and (C). In general, our GP-based estimates have smaller SRE and larger ENV than the EM-based estimates and hence, the MRW is usually wider in the GP-based estimates, accurately reflecting the uncertainty of the estimates. As expected, increasing the number of loci (m) generally decreases the width of the confidence and credible intervals of our estimates (MRW). Although this is generally true for EM estimates as well, EM estimates have very low coverage of the truth (MRE in Table 4) when the number of loci increases.

**Figure 6:**
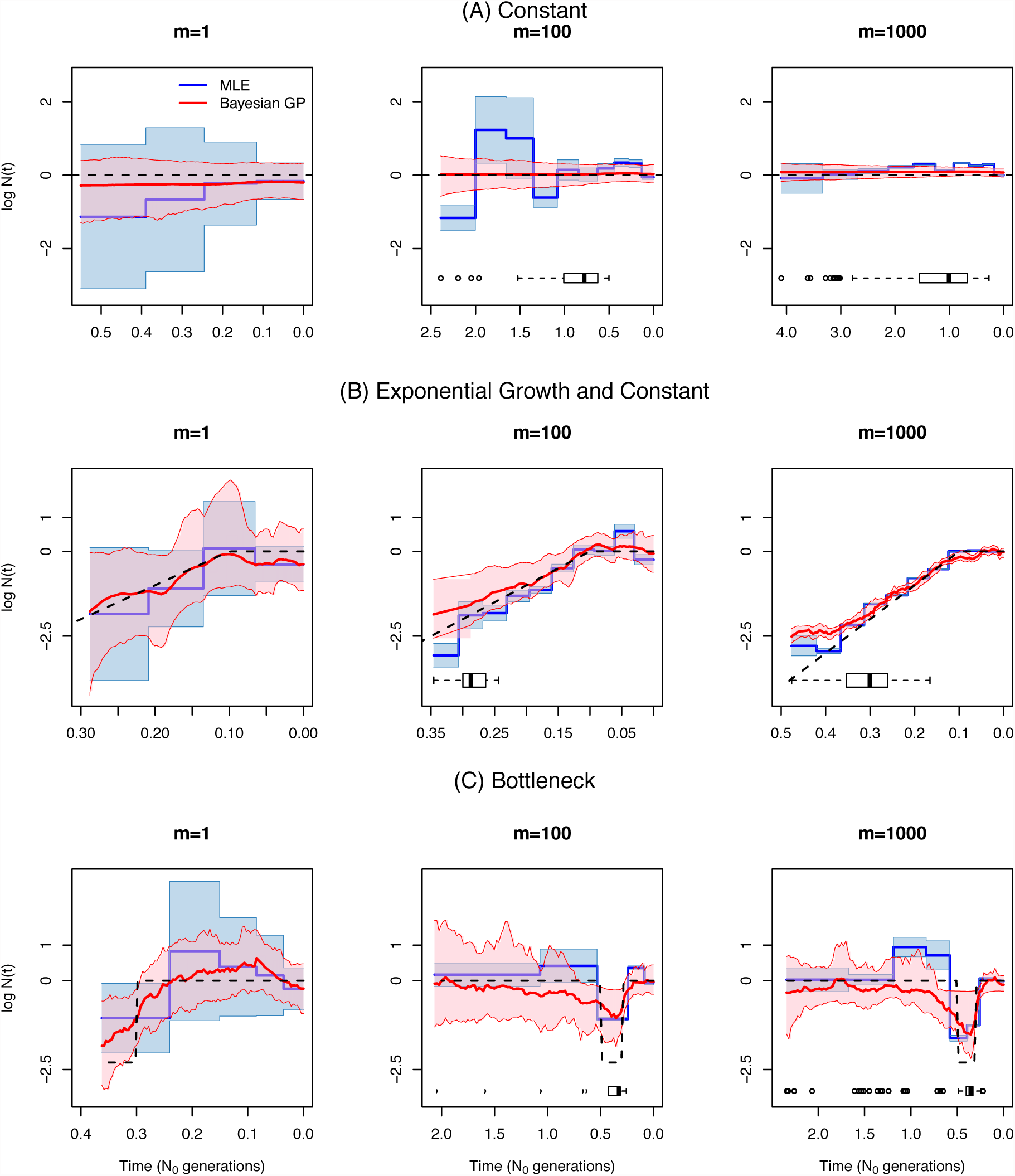
Inference of population size trajectories *N*(*t*) for *n* = 20. (A) Simulated data under constant population size, (B) exponential and constant trajectory, and (C) a bottleneck. We show estimates from *m* = 1 genealogy, *m* = 100 local genealogies and *m* = 1000 local genealogies. We show the true trajectories as dashed lines, blue lines and light blue shaded areas represent EM point estimates and 95% confidence areas, and red lines and pink shaded areas represent Bayesian GP posterior medians and 95% BCIs. Boxplots of the TMRCA are shown at the bottom of each plot.

**Figure 7:**
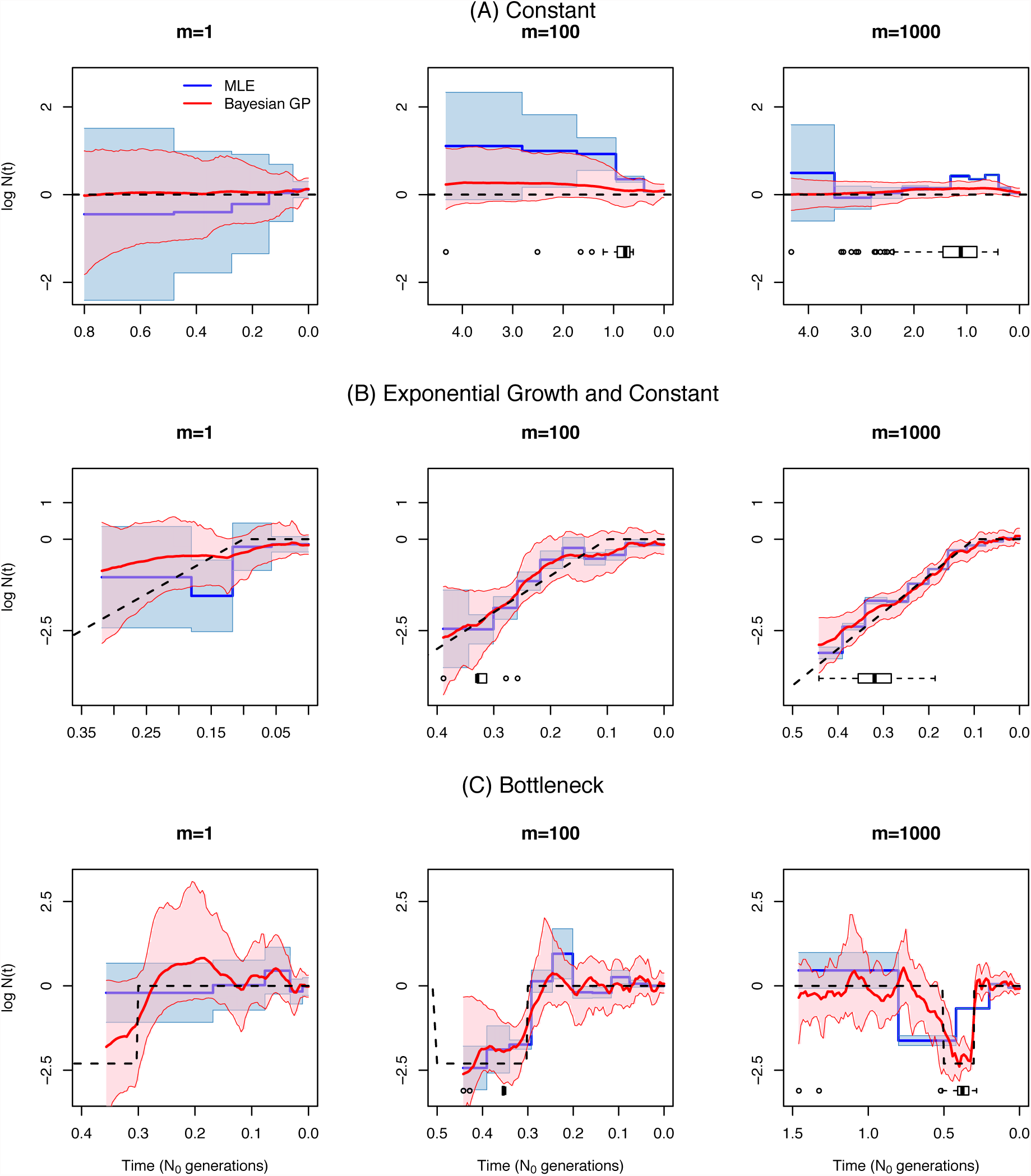
Inference of population size trajectories *N*(*t*) for *n* = 100. (A) Simulated data under constant population size, (B) exponential and constant trajectory, and (C) bottleneck. We show estimates from *m* = 1 genealogy, *m* = 100 local genealogies and *m* = 1000 local genealogies. We show the true trajectories as dashed lines, blue lines and light blue shaded areas represent EM point estimates and 95% confidence areas, and red lines and pink shaded areas represent Bayesian GP posterior medians and 95% BCIs. Boxplots of the TMRCA are shown at the bottom of each plot.

### 4.3 Sampling more individuals versus sequencing more loci

Figures 5-7 show our estimates for *n* = 2, 20 and 100 sampled individuals across varying numbers of loci. Since performance of EM estimates depends strongly on the definition of the grid, we base what follows on the Bayesian GP estimates. We find that increasing the number of loci, decreases uncertainty of our estimates and allows us to infer *N*(*t*) further back in time. Increasing the number of samples does not necessarily increase the performance of our GP estimates. For example, under the bottleneck scenario, we are able to detect the bottleneck phase fairly accurately even for two samples with *m* = 1000 local genealogies. Increasing the number of samples to *n* = 20 and *n* = 100 does not improve estimation of the features of the bottleneck. This is because most TMRCAs observed under the bottleneck scenario occur during the bottleneck (Figures 5,6 and 7), regardless of the number of individuals sampled. In contrast, in our exponential growth example, increasing the number of samples from *n* = 2 to *n* = 100 improves accuracy (point estimates are closer to the truth, see SRE in Tables 4A-C) and credible intervals cover the truth completely (ENV of 100%).

### 4.4 Sequential Tajima’s genealogies are sufficient statistics under the SMC ′

Under the SMC′ process, marginally at each locus along the chromosome, a local genealogy is a realization of Kingman’s n-coalescent (Kingman 1982), a continuous-time Markov chain taking its values in the set *𝒦*_*n*_ of partitions of the label set {1, 2,…, *n*}. A local genealogy *g* of *n* individuals includes labeled topology *𝒦*_*n*_ and coalescent times *t* = (*t*_*n*_,…,*t*_2_). The state space of a local genealogy is then 𝒢 = *𝒦*_*n*_ ⊗ R^+*n−*1^, and the cardinality of the set *𝒦*_n_ is *n*!(*n* − 1)!/2^*n*−1^. However, only the set of ordered coalescent times carry information about *N*(*t*). For a single locus, the set coalescent times are sufficient statistics for inferring *N*(*t*) (proof is in the Appendix). A natural question that follows is whether the coalescent times corresponding to the set of local genealogies are sufficient statistics for inferring *N*(*t*) under the SMC′ model. We find that the sufficient statistics for inferring *N*(*t*) under the SMC′ model, are the coalescent times, when taken together with local ranked tree shapes. For a single locus, the set of coalescent times together with the ranked tree shape correspond to a realization of Tajima’s n-coalescent. Tajima’s n-coalescent (Tajima 1983) is a continuous-time Markov chain taking its values in the set H*_n_* of ranked tree shapes also called histories, evolutionary relationships or vintaged and sized coalescent (Sainudiin et al. 2014). The state space of Tajima’s local genealogy is then *𝒢^T^* = *ℋ_n_* ⊗ *ℝ*^+*n*-1^, and the cardinality of the set *ℝ_n_* corresponds to the sequence of Euler zigzag numbers whose first ten elements are 1, 1, 1, 2, 5, 16, 61, 272, 1385, 7936 (Disanto and Wiehe 2013). The probability of getting a particular type of ranked tree shape *H_n_* of *n* samples (Tajima 1983) is given by

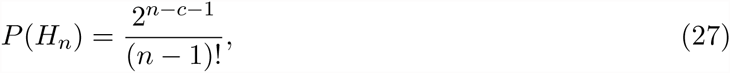

where *c* is the number of *cherries*, defined as branching events that lead to exactly two leaves.

In the Methods section, we defined transition densities in terms of coalescent times and *F_i,j_* quantities. The set of all *F_i,j_* quantities from a local genealogy form a triangular matrix: *F*-matrix. In the Appendix, we show that (*i*) *F*-matrices are in bijection with ranked tree shapes and (*ii*) the set of local Tajima’s genealogies are sufficient statistics for inferring *N*(*t*) under the SMC′ model. These observations are crucial for inferring *N*(*t*) from sequence data directly. Coalescent-based inference from sequence data rely on marginalization over the hidden state space of genealogies. In the Appendix, we show that the state space needed is the space of local Tajima’s genealogies, as opposed to the space of local Kingman’s genealogies. For *n* = 10 sequences, there are 2, 571, 912, 000 possible labeled topologies while only 7, 936 possible ranked tree shapes.

### 4.5 Application to human data

We applied our method to a 2-Mb region on chromosome 1 (187,500,000-189,500,000) with no genes from five Yorubans from Ibadan, Nigeria (YRI) and five Utah residents of central European descent (CEU) from the 1000 Genomes pilot project (1000 Genomes Project Consortium 2012) and previously analyzed for the same purpose (Sheehan et al. 2013). We used *ARGweaver* (Rasmussen et al. 2014) to obtain a sample path of local genealogies for the two populations (YRI and CEU). The parameters used are 200 change points, a mutation rate of *μ* = 1.26×10^−8^ and a recombination rate of *ρ* = 1.6 × 10^-8^ (Rasmussen et al. 2014, details regarding parameters used can be found in Supplementary Information). We note that *ARGweaver* assumes the SMC process and our method assumes the SMC′ process. Moreover, our inference is based on a single sample of the SMC process with known pruning times. Our *ARGweaver* set of local genealogies are discretized at 200 time points and our GP-based inference is influenced by this discretization. In Figure 8 we show our estimates of past Yoruban (in blue) and European population sizes (in green). The two population size trajectories experience a series of bottlenecks and overlap until about 100 YKA, assuming a diploid reference population size of *N*_0_=10,000 and a generation time of 25 years. In Figure 8 we recover an out-of-Africa bootleneck that starts about 100 KYA and ends about 30 KYA in the European population. These results are consistent with previously published results (Li and Durbin 2011; Gronau et al. 2011; Rasmussen et al. 2011; Sheehan et al. 2013; Schiffels and Durbin 2014). In Supplementary Information Figure S4 we show the estimates of *logN*(*t*) instead of *logN*(*t*) and time measured in units of *N*_0_ generations (same scaling as with simulations in Figures 5-7).

**Figure 8:**
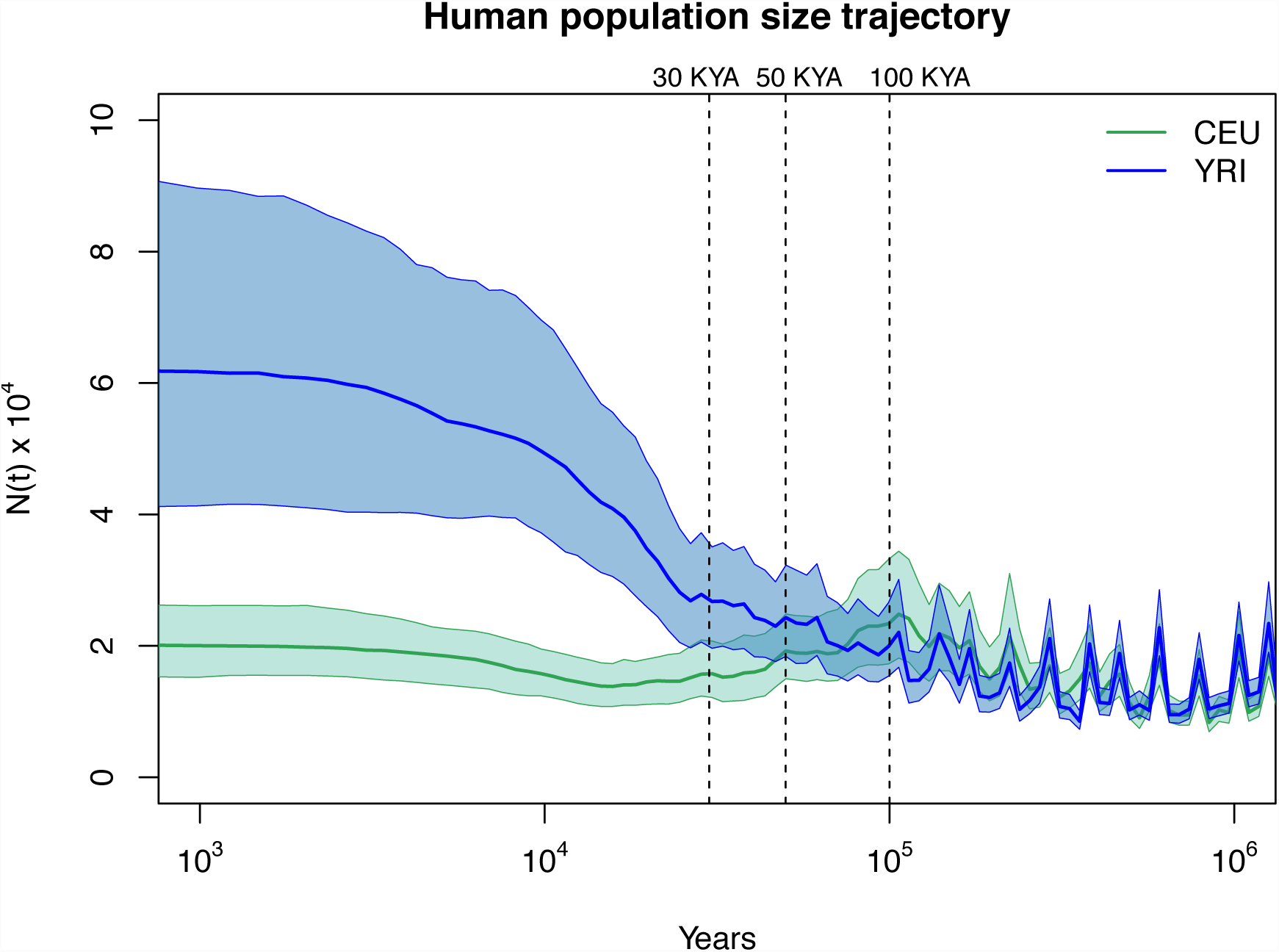
Inference of human population size trajectories *N*(*t*) for *n* = 10. Green solid line and green shaded areas represent the posterior median and 95% BCI for European population (CEU) and blue solid line and blue shaded areas represent the posterior median and 95% BCI for Yoruban population (YRI). Time is measured in years in the past assuming a generation length of 25 years and a reference diploid population of 10,000 individuals. The x-axis is log transformed.

## Discussion

In this paper, we propose a Gaussian-process based Bayesian nonparametric method for estimating effective population size trajectories *N*(*t*) from a sequence of local genealogies, accounting for recombination. Under a variety of simulated demographic scenarios and sampling designs, our method recovers the truth with better precision and accuracy than a maximum likelihood approach (Figures 5-7). We apply our method to genealogies estimated from human genomic data *ARGweaver* (Rasmussen et al. 2014) and conduct inference of the human population size trajectory for European and African populations; this application to real data recover the known features of the out of Africa bottleneck (Figure 8).

Several recent approaches have emerged for inference of population size trajectories from multiple whole-genome sequences using the sequentially Markov coalescent (SMC) (Li and Durbin 2011; Sheehan et al. 2013; Schiffels and Durbin 2014). However, current SMC-based methods rely on maximum likelihood inference (EM) of both a discretized parameter space and a discretized state space in order to gain computational tractability, and incur the costs of reduced accuracy and biased estimates. Although in principle the EM approach and the Bayesian nonparametric approach approximate *N*(*t*) similarly — by either a piece-wise constant or a piece-wise linear function — the Bayesian nonparametric approach is not affected by increasing the number of parameters (or change points) in the estimation of *N*(*t*). For comparison with existing methods, we implemented an EM approach to infer population size trajectories from a sequence of local genealogies and we note that increasing the number of loci may actually increase the bias of the EM estimates (Figures 5-7). For example, in simulation, our EM approach incorrectly detects the initial period of the simulated bottleneck (around 0.8*N*_0_ instead of 0.5*N*_0_ generations ago) with narrow confidence intervals (Figure 7C).

There are many advantages to using Bayesian GP over EM for inference of population size trajectories. Similar to Palacios and Minin’s (2013) approach to inference from a single genealogy, we a *priori* assume that *N*(*t*) follows a log Brownian Motion process. This allows us to model *N*(*t*) as a continuous positive function. The main advantage of using a Brownian Motion process is that its inverse covariance function is a sparse matrix that allows for fast computations. Since the likelihood function involves integration over *N*(*t*), this integral is approximated by the Riemann sum over a regular grid of points. The finer the grid is, the better the approximation. We find that our method performs well for inferring *N*(*t*) at 100 change points in all our examples and, more importantly, results are not sensitive to the number of change points used in the analysis (Figure 4). Our Bayesian approach relies on MCMC for inference from the posterior distribution of model parameters. Because population sizes at different grid points are correlated, we adapt the recently developed MCMC technique Split Hamiltonian Monte Carlo (splitHMC) for jointly sampling all model parameters (Shahbaba et al. 2014; Lan et al. 2015). splitHMC is a Metropolis sampling algorithm that efficiently proposes states that are distant from current states with high acceptance rates. It has been shown to be more efficient in inferring *N*(*t*) from a single genealogy than elliptical slice sampling or regular Hamiltonian Monte Carlo sampling(Lan et al. 2015). However, splitHMC relies on calculating the score function at every single iteration. Because pruning time in each local genealogy is unknown, we calculate the score function via Fisher’s formula.

In simulations, we find that our algorithm scales well with hundreds of individuals; our computational bottleneck is in the number of local genealogies. We envision that extending the current methodology to inference from sequence data directly will require a strategy for sampling shorter genomic segments. This would be a probabilistic alternative to arbitrarily choosing segment lengths (Sheehan et al. 2013; Rasmussen et al. 2014).

Under the SMC model, every recombination event along the genome translates to a new coalescent event for the sample under study, so increasing the number of loci results in more realizations of the coalescent process. The longer the segments are and the larger the number of samples taken, the greater the chance of observing variation due to recombination. This fact makes it hard to define a sampling strategy: longer genomes or larger sample sizes? We show that increasing the number of local genealogies improves precision of our Bayesian GP estimates (Figures 5-7). However, resolution into the past from contemporaneous sequences highly depends on the actual population size trajectory *N*(*t*).

We use *ARGweaver* (Rasmussen et al. 2014) to generate two samples of contiguous local genealogies corresponding to a 2-Mb region of chromosome 1 for five Europeans (CEU) and five Africans (YRI) from the 1000 Genomes Project; this genomic region is free of genes and was also analyzed in Sheehan et al. (2013). Taking these two samples of local genealogies as our data (4186 local genealogies for CEU and 6247 local genealogies for YRI), we were able to use our Bayesian GP method to infer Yoruban and and European effective population size trajectories (Figure 8). We find an out-of-Africa bottleneck that began ∼ 100 KYA and ended ∼ 30 KYA in the European population consistent with Li and Durbin (2011); Rasmussen et al. (2011); Gronau et al. (2011); Sheehan et al. (2013) and Schiffels and Durbin (2014). We note that our estimates are based on a single sample of local genealogies and thus ignore genealogical uncertainty. Moreover, we generated our data from the posterior distribution of local genealogies using *ARGweaver* at 200 time intervals so our GP-based approach cannot fully detect sudden changes that may occur between the discretized times. In addition, *ARGweaver* assumes an SMC prior model on local genealogies and our GP-based method assumes the SMC′ process; the lack of invisible recombination events in *ARGweaver* ‘s genealogies will bias inference.

The natural next extension for our method presented in this study is to infer *N*(*t*) from sequence data directly and not from the set of local genealogies. Our MCMC approach allows us extend the current methodology in a Bayesian hierarchical framework where the SMC′ process would be used as a prior distribution over local genealogies. The work we present here suggests a combination of *ARGweaver* accommodating SMC′ and GP priors would result in an efficient method for inferring population size trajectories from sequence data directly. In addition, our model can be easily modified to model a variable recombination rate along chromosomal segments and to jointly infer variable recombination rates and *N*(*t*).

Finally, we show that, under the SMC′ model, local ranked tree shapes and coalescent times correspond to a set of local Tajima’s genealogies; these Tajima’s genealogies are the sufficient statistics for inferring *N*(*t*). Under the SMC′ model, the state space needed for inferring population size trajectories from sequence data is that of a sequence of local Tajima’s genealogies. This lumping, or reduction of the original SMC′ process, will allow more efficient inference from sequence data directly.

Current methods for inferring population size trajectories make tradeoffs to analyze whole genomes that limit both biological understanding of sudden population size changes and the ability to test hypotheses regarding population size changes. This work represents a critical set of theoretical results that lay the groundwork for efficient estimation of detailed histories from sequence data with measures of uncertainty.

## Acknowledgements

We thank Amandine Veber for her valuable suggestions and comments. We thank Shiwei Lan for his suggestions for speeding up the MCMC sampling scheme and Sara Sheehan and Melissa J. Hubisz for helpful discussions. J.A.P. acknowledges scholarship from CONACyT Mexico to pursue her research work. This research is supported in part by NSF CAREER Award DBI-1452622 (to S.R.). S.R. is a Pew Scholar in the Biomedical Sciences, supported by The Pew Charitable Trusts, and an Alfred P. Sloan Research Fellow.

## Appendix A

### Discretization

For both our Bayesian method and our EM method, we assume that *N*(*t*) is a piece-wise linear (or piece-wise constant) function with *d* change points. Let 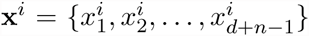 be the ordered set of time points corresponding to the change points *x*_1_,…,*x*_*d*_ and the coalescent time points **t**^i^ of local genealogy *i*. Then, we calculate all the factors needed for the observed data likelihood (Equation 13) and the complete data likelihood (Equation 20).

Let 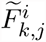 denote the discretized version of *F ^i^* that represents the number of branches in *g*_*i*_ that do not coalesce with any other branch in the time interval 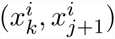. Note that the indices here are in increasing order, *k ≤ j*. Similarly, let 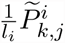 denote the probability that *U*_*i*_ (the time along genealogy *i*), occurs in 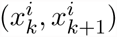 and the self-coalescing event occurs at time 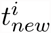 in 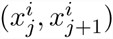.That is

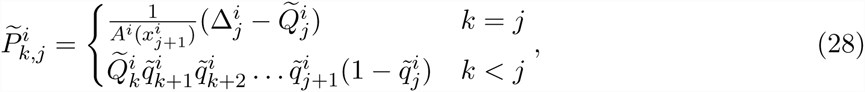

where

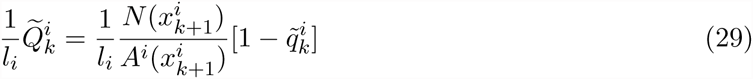

is the joint probability of pruning time 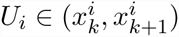 and not coalescing back to the same branch in the time interval 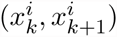, and

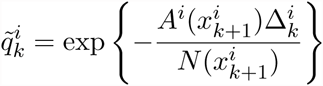

### Expectation-Maximization Algorithm

**E-step:** Equations 22 and 23 show that for the E-step, the only expectations we need are 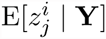 and 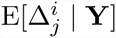. We compute these expression as follows:

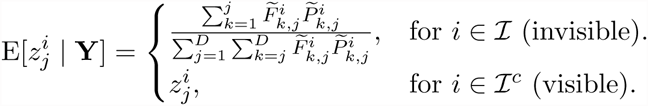

For *i* ∈ ℐ

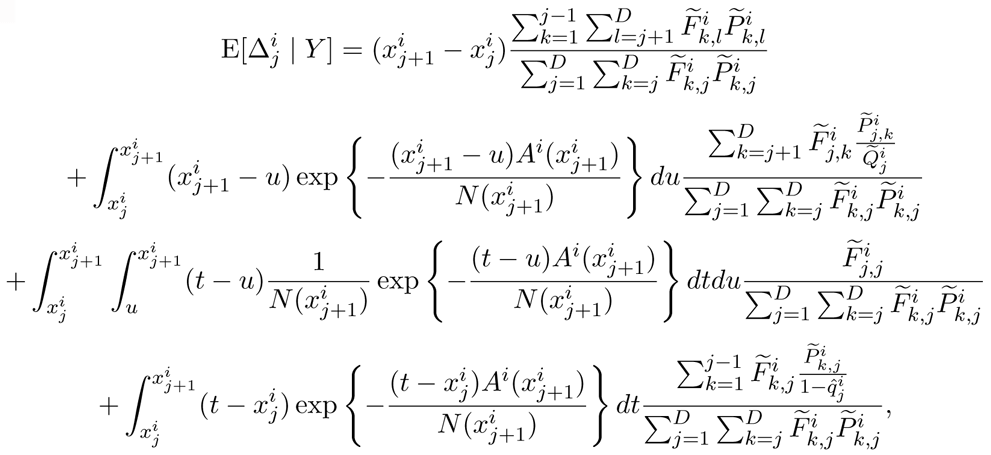

and for *i* ∈ ℐ^c^, let

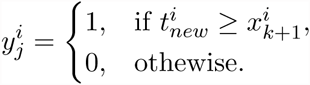

Then

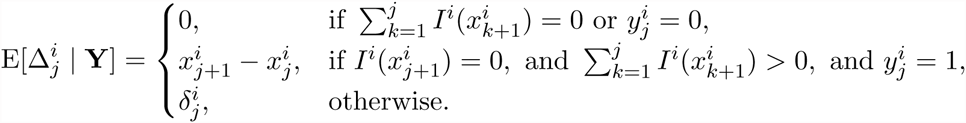

where

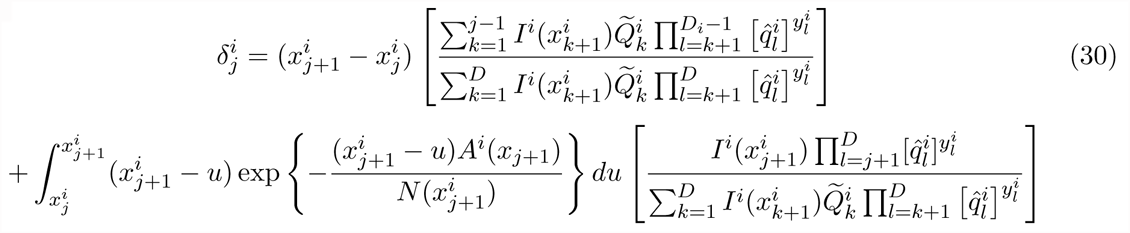

**M-step.** Now, for the kth iteration of the algorithm and maximizing the complete data loglikelihood (Equation 20) we have

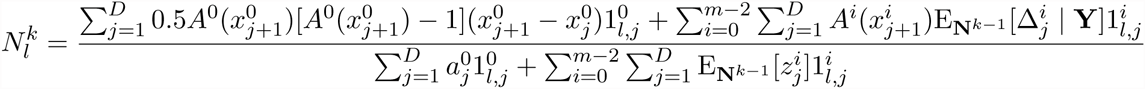

where

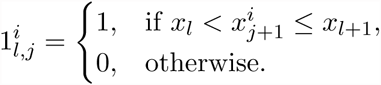

is an indicator function that takes the value of 1 when (*x*_*l*_, *x*_*xl*+1_) covers the interval 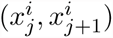.

### Observed score function for Split Hamiltonian Monte Carlo

Our Bayesian approach relies on Split Hamiltonian Monte Carlo (splitHMC) to sample from the posterior distribution of model parameters. This method requires the calculation of the observed score function. We use Fisher’s identity and calculate the observed score function as the conditional expected complete score function. The *l*th element of ∇ℒ_*obs*_ is

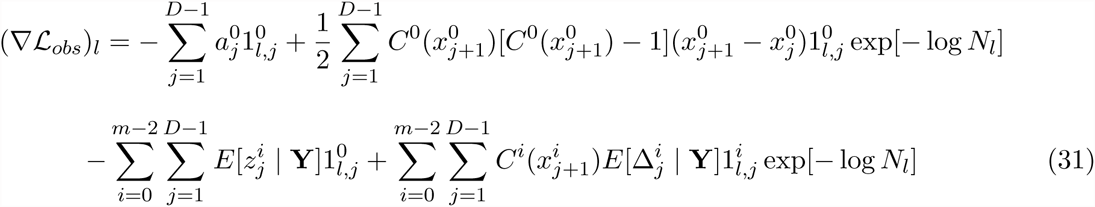

### Fisher Information Calculation

The calculation of the Fisher information needed to estimate confidence intervals of a piece-wise constant trajectory of population sizes, requires the following expected values:

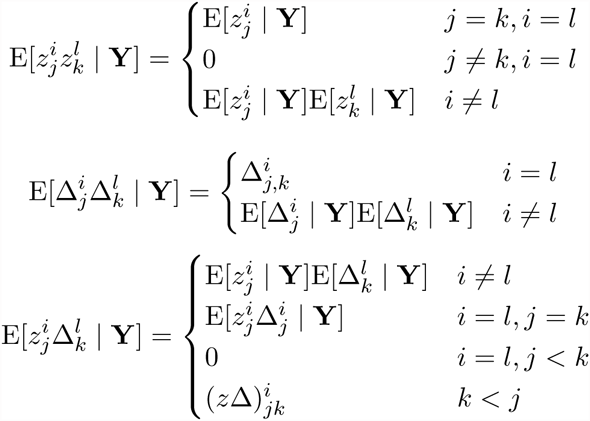

For *k* < *j* and *i* ∈ ℐ

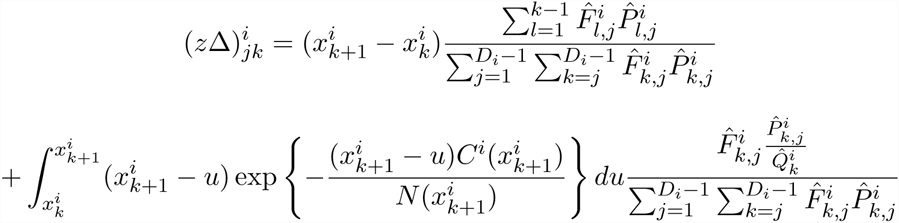

and for *k* < *j* and *i* ∈ ℐ^c^

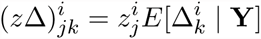

For *j* < *k* and *i* ∈ ℐ

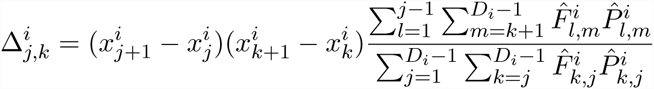

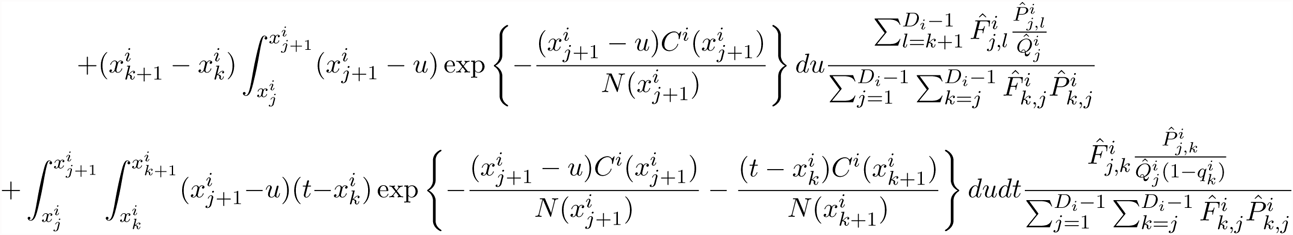

and

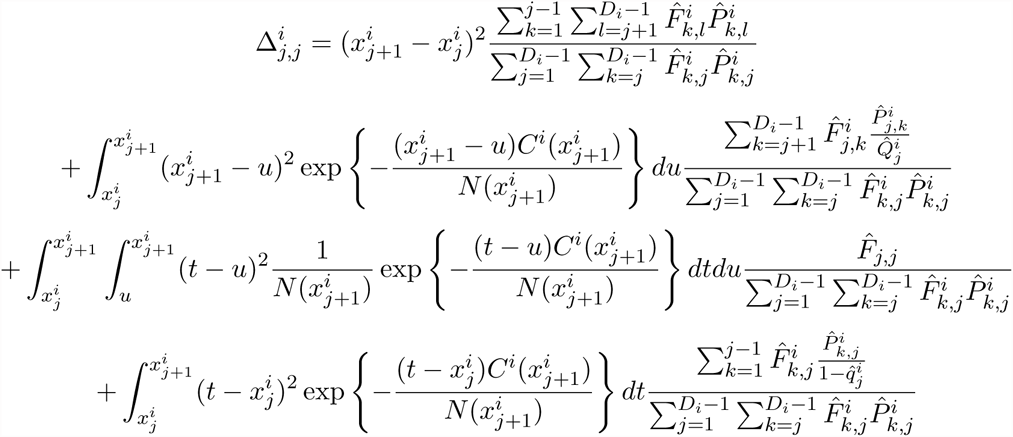

For *i ∈ ℐ^c^* and *j* < *k*

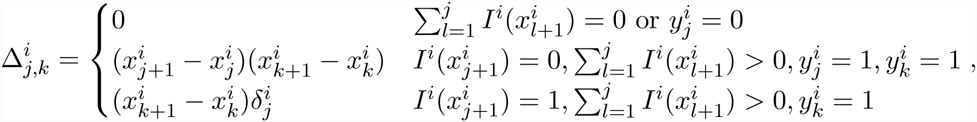

where 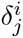 is as defined in Equation 30, and

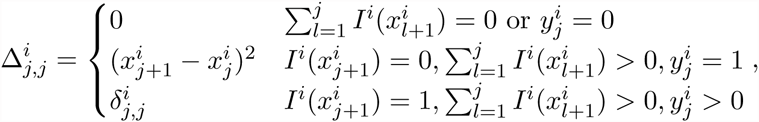

where

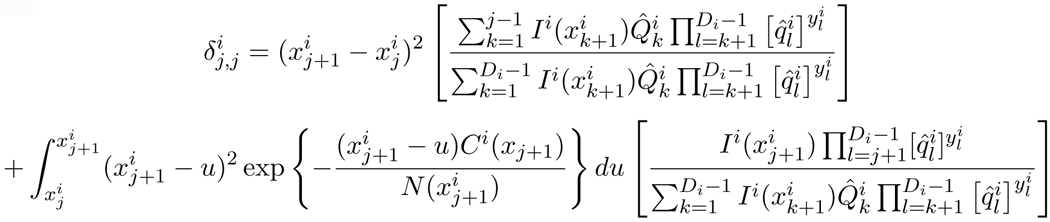

and

For *i ∈ ℐ*

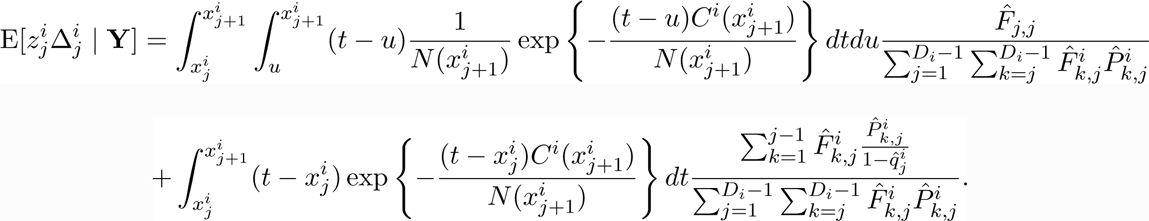

and for *i* ∈ *ℐ^c^*

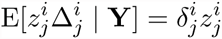

The gradient vector of the complete data log-likelihood has *l*th element

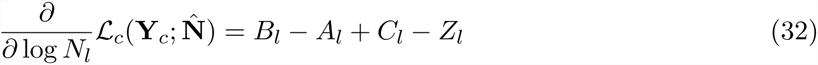

With

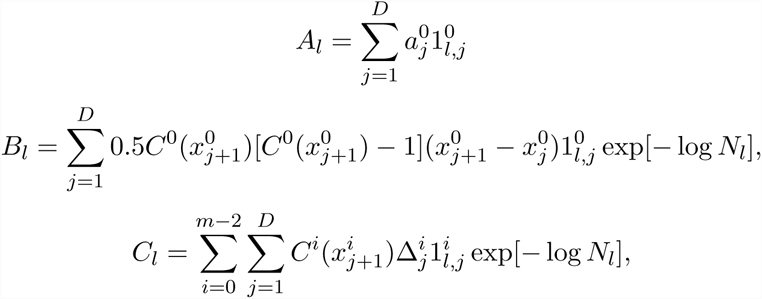

and

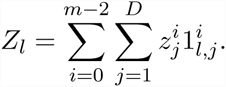

Next, differentiating Equation (32), we have 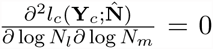 = 0 for all *l* ≠ *m*, so the Hessian is a diagonal matrix with (*l, l*)th element

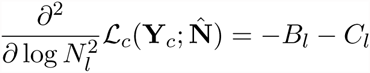

and

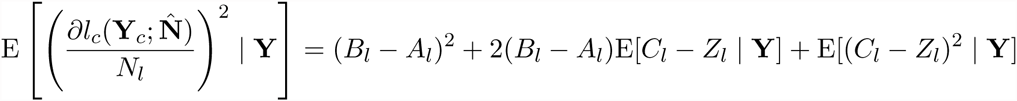

where

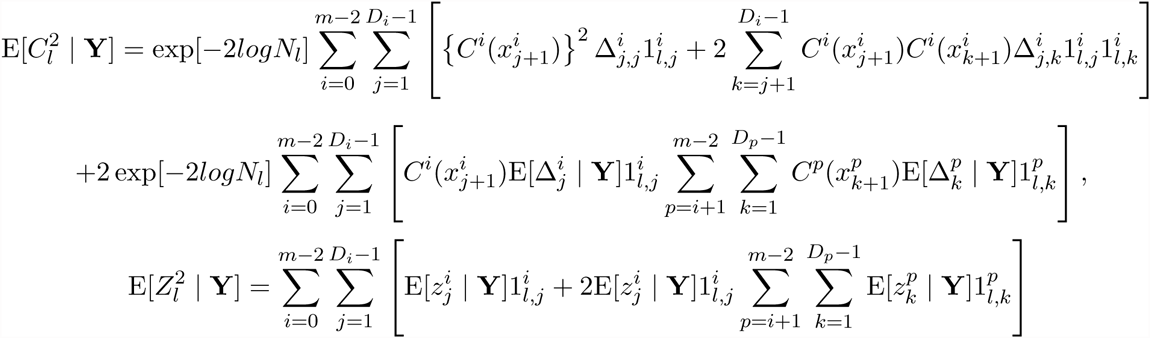

and

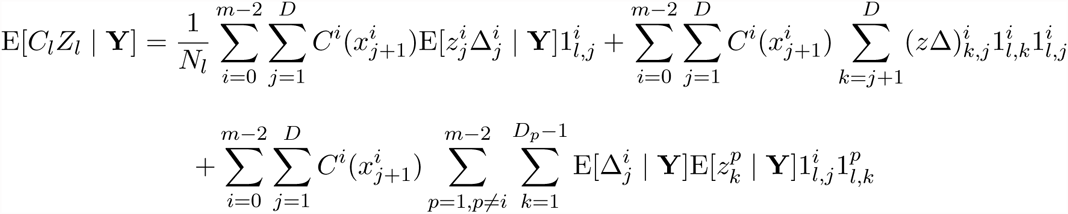

Also,

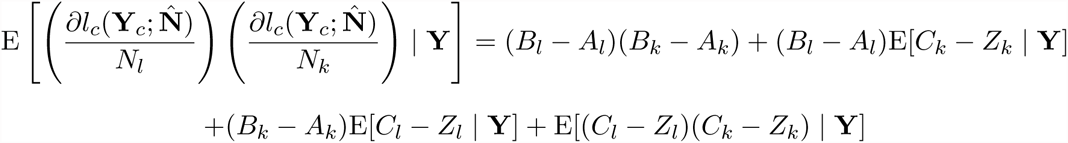

where

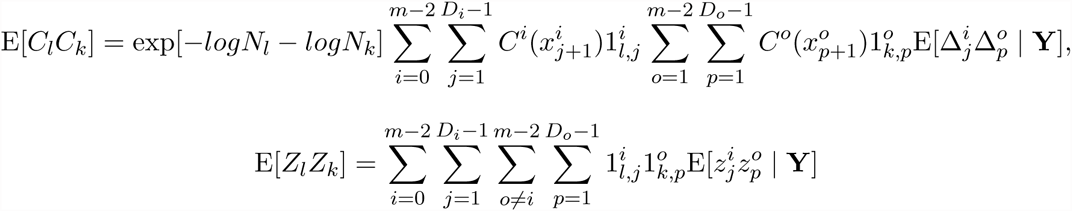

and for *l* < *0*

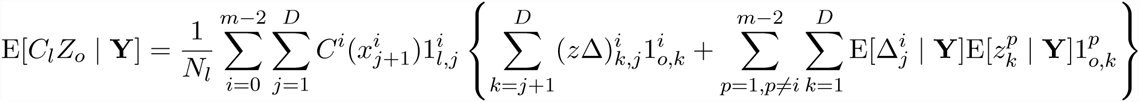

### Sufficient statistics under SMC′

#### Proposition 1.

*For a single locus, the set of coalescent times are sufficient statistics for inferring *N*(*t*).*

*Proof.* This can be proved using the factorization theorem. The marginal density of a local genealogy (Equation 3) has a unique factor that depends on *N*(*t*) and *g* only through *t*_*n*_,…,*t*_2_. The values of *A*(*t*) are induced by the natural order of the coalescent times.

Let *F* denote a lower triangular matrix of size *n* × *n* with the *F_i,j_* entry the number of lineages that do not coalesce in the time interval (*t*_*i*+1_, *t*_*j*_), as defined in the methods section and with the following properties:

1. *F*_*i*,1_ = 0 for all *i* = 1,…,*n* (The first column contains 0s for completion)
2. *F*_*i,j*_ = 0 for all *j* > *i* (Lower triangular matrix)
3. *F*_*i,i*_ = *i* for all *i* ≥ 2 (The diagonal corresponds to the number of lineages at each intercoalescent interval)
4. *F*_*i,i*−1_ = *i* − 2 for all *i* ≥ 2 (At each intercoalescent interval, we loose two free lineages, so the second diagonal correspond to the number of lineages minus two)
5. For *j* < *n* − 1, the last row of *F* is defined according to:

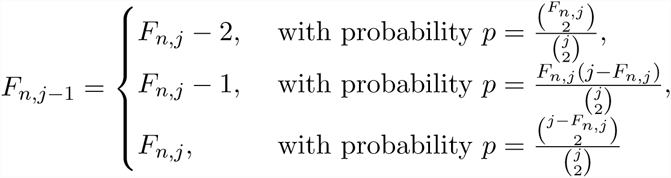
6. Let *c* denote the number of cherries, then

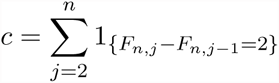
7. For *i* < *n* and *j* < *i* − 1, if *F*_*n,j−*1_ = *F*_*n,j*_ − 2, then *F*_*i,j*−1_ = *F_i,j_* − 2.
8. Let *v_i_* denote the set of lineages in the intercoalescent interval (*t*_*i*_, *t*_*i-*1_) with direct descendant internal nodes. The lineage labels correspond to the label of the coalescent time, when the direct descendant internal node was created. That is, the lineage created at *t*_*n*_ has label *n*: *v_n_* = {*n*}; the lineage created at *t*_*i*_ has label *i*. Let |v*_i_*| denote the size of the set v*_i_*. Note that 1≤ |*v_i_*| ≤*c* and

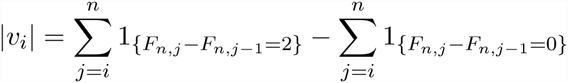
9. For *i* < *n* and *j* < *i* − 1, if *F*_*n,j−*1_ = *F*_*n,j*_ − 1, then at time *t*_*j*_, there is a coalescence between a singleton and a lineage in the set *v*_*j*_. Let *a*_*j*_ be the lineage selected uniformly at random

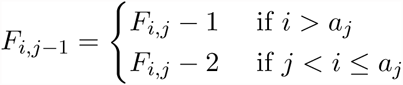
10. For *i* < *n* and *j* < *i* – 1, if *F*_*n,j-*1_ = *F_n,j_*, then at time *t*_*j*_, there is a coalescence between two lineages 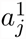 and 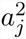 from the set v*_j_*. Let 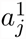 denote the minimum and 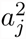 the maximum of the two lineages selected, then

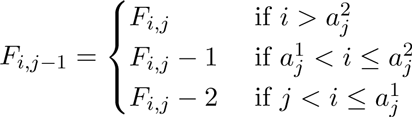

We show the correspondence between a ranked tree shape and the F-matrix in the example of Figure A1. The first row and the first column are set to 0, the first two diagonals are known with probability 1: *F_i,i_* = *i* and *F*_*i,i-*1_ = *F_i,i_* − 2 for *i* > 1. In our example, *n* = 5 and so, the first diagonal corresponds to (0, 2, 3, 4, 5) and the second diagonal corresponds to (0, 1, 2, 3). The last row *F*_5_, contains 0, followed by the number of branches that do not coalesce in the time intervals (*t*_6_, *t*_2_), (*t*_6_, *t*_3_), (*t*_6_, *t*_4_) and (*t*_6_, *t*_5_) corresponding to (0, 0, 2, 3, 5).

**Figure A1:**
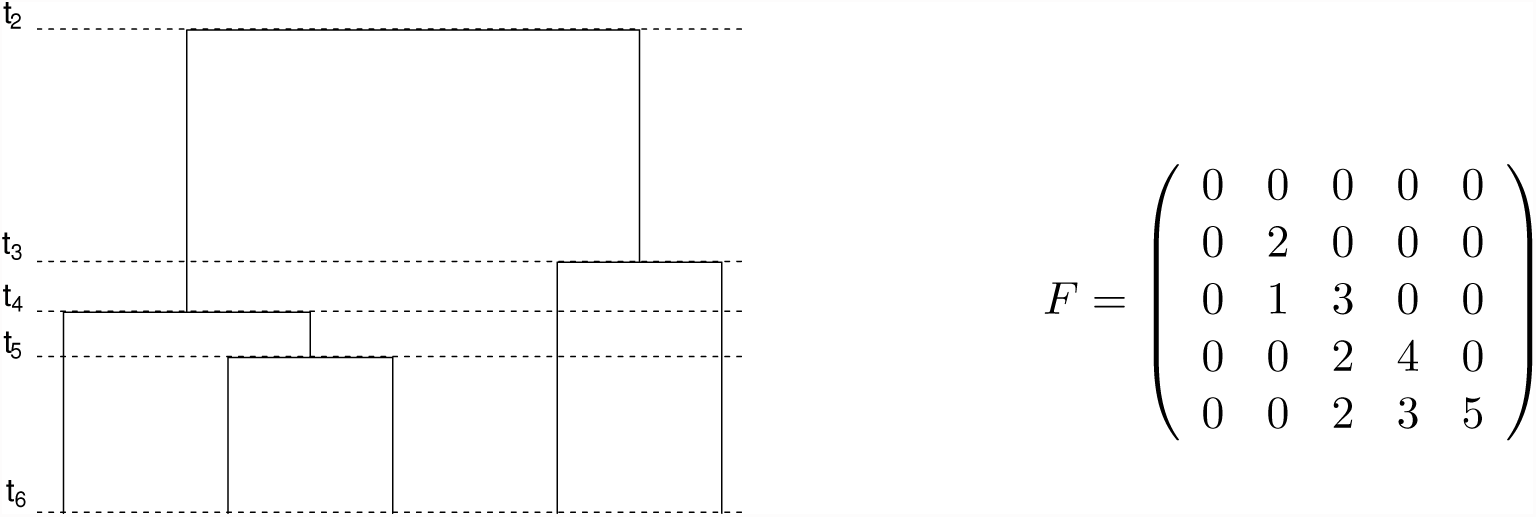
Ranked tree shape for *n* = 6

#### Proposition 2.

*There is a bijection between the set of ranked tree shapes* ℋ*_n_ and* ℱ*, the set of *F*-matrices.*

*Proof.* The probability of the *F* matrix can be expressed as the product of the conditional probabilities of the columns of the *F* matrix, that is:

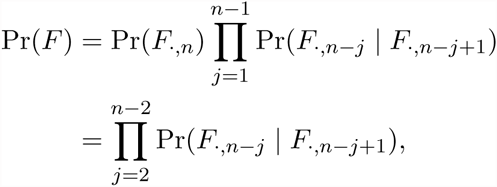

since the first and last column of *F* are known with probability 1. Note *F*_*.,j*_ represents the *j*-th column vector of the *F* matrix.

Let *d*_*i*_ = *F*_*n,i*_ − *F*_*n,i-*1_ for *i* = 3,…,*n*, and *d*_2_ = *F*_*n,*2_ then

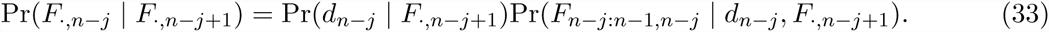

That is, the conditional probability of the (*n − j*)th column of *F* given the (*n − j* + 1)th column of *F* is the product of the conditional probability of the last element of the (*n − j*)th column and the conditional probability of the rest of the (*n − j*)th column. When d*_n-j_* = 2 the rest of the column is known with probability 1 (property 7 of the *F*-matrix). When d*_n-j_* = 1, the rest of the *n − j*th column has probability 1/|*v*_*n-j*+1_| (property 9 of the *F*-matrix) and when d*_n-j_* = 2, the rest of the *n* − *j*th column has probability 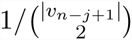 (property 10 of the *F*-matrix). Then re-writing Equation 33, we have

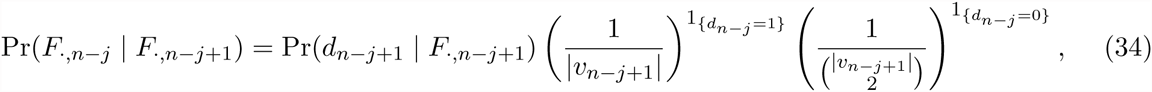

since 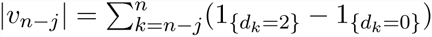, and 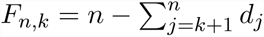, then

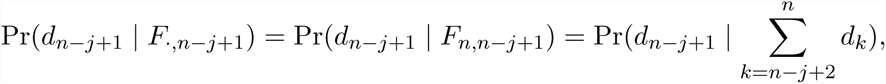

and

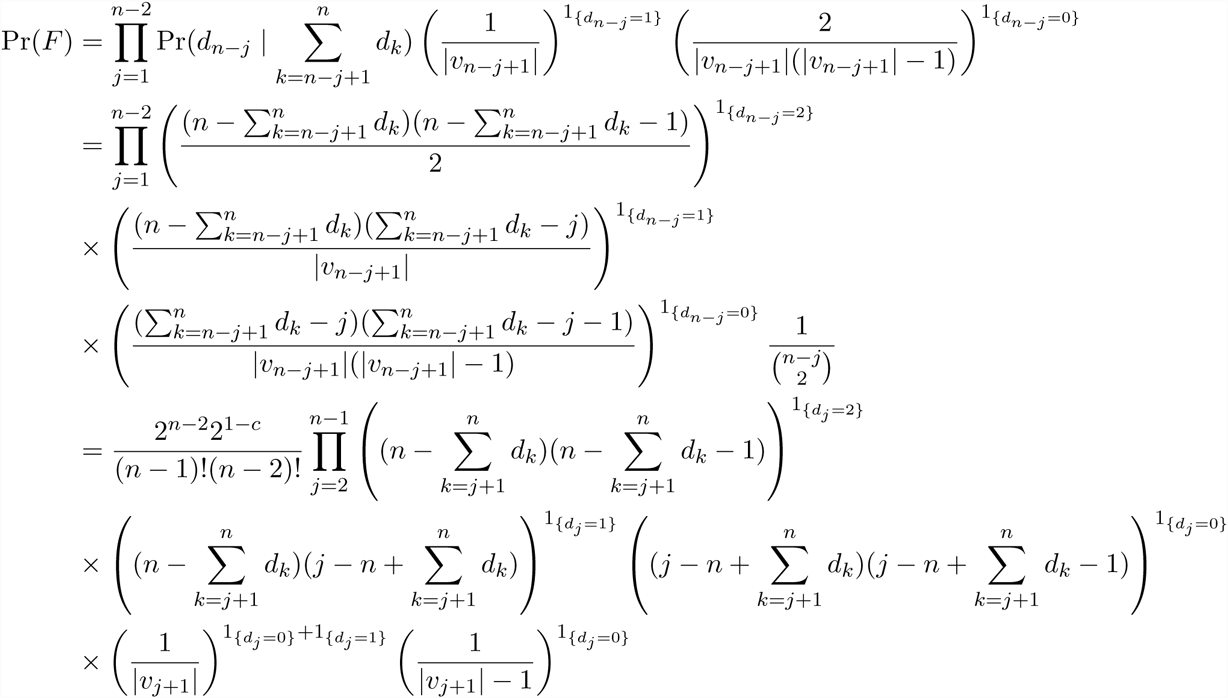

Since *d_n_* = 2 and |*v_n_*| = 1, for *j* = *n* − 1, then *d*_*n-*1_ is either 1 or 2, then

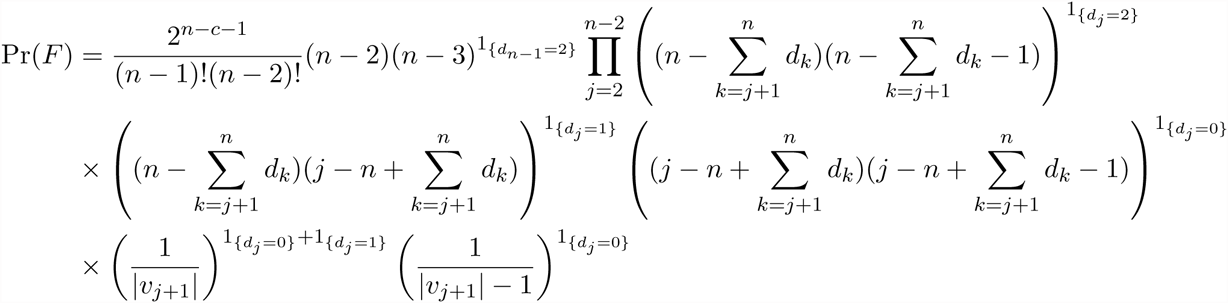

If we continue expanding the expressions, we get:

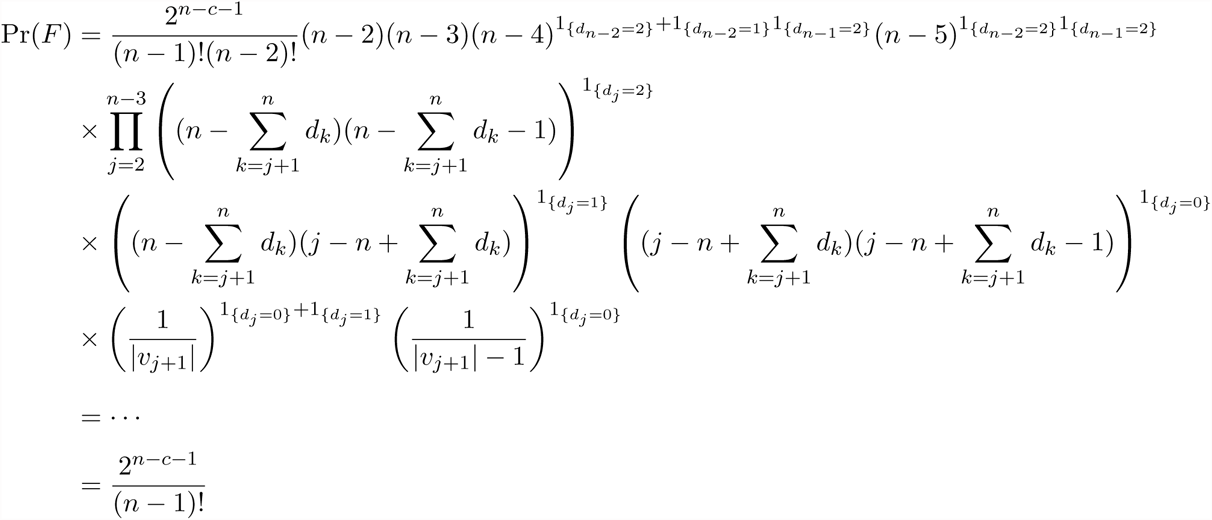

Note that the entries of the *F* matrix correspond to the same quantity needed to express the transition density of an invisible event (Equation 11). We claim that the sequence of coalescent times sets **t**^0^, **t**^1^,…,**t**^*m-*1^ and *F* ^0^,F ^1^,…,*F*^*m-*1^ matrices corresponding to the ranked tree shapes of local genealogies *g*_0_, *g*_1_,…,*g*_*m-*1_ are sufficient statistics to infer *N*(*t*) under the SMC′ process. We prove this through the following propositions.

#### Proposition 3.

*The probability density of Tajima’s genealogy is proportional, up to a combinatorial factor, to the probability density of Kingman’s genealogy.*

*Proof.*

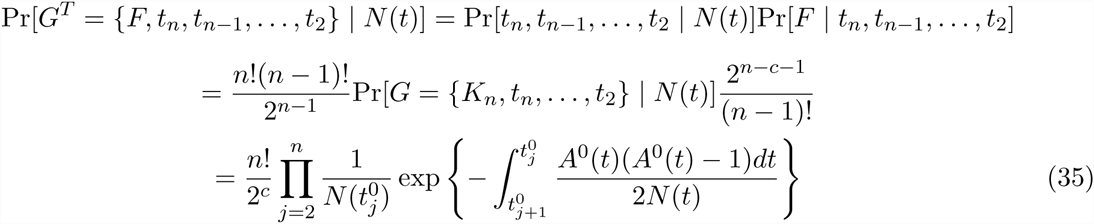

#### Proposition 4.

*The marginal visible transition density from a local Kingman’s genealogy g*_*i-*1_ *to G_i_ is proportional to the marginal visible transition density from the corresponding local Tajima’s genealogy* 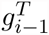 to 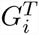

*Proof.* When the labeled topology of *g*_*i-*1_ is the same as the labeled topology of *g*_*i*_, then a transition from *g*_*i-*1_ to *g*_*i*_ contains the same information about pruning location as a transition from 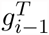 to 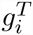 (Supplementary Information, Figures S1A and S2D). In fact, the *I*^*i*-1^(*t*) function defined in section 2.1.2 (Equation 8) can be defined in terms of the *F ^i^*-matrix and the coalescent times **t**^*i*-1^ and **t***^i^*. In this case, for some *j* ∈ 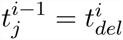 and 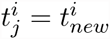. Then

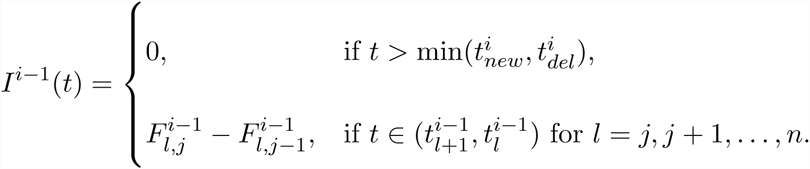

Hence, if *K*_*i-*1_ = *K*_*i*_, the labeled topologies of *g*_*i-*1_ and *g*_*i*_, then

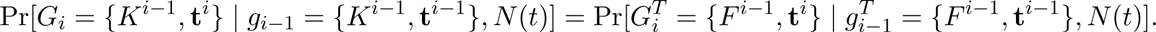

When the labeled topologies of *g*_*i-*1_ and *g*_*i*_ are different, but the children of 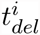and the children of 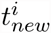 are the same, we cannot exactly identify the pruning branch and the new coalescing branch (Supplementary Information, Figure S1B) and then a transition from *g*_*i-*1_ to *g*_*i*_ contains the same information about pruning location as a transition from 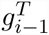 to 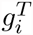. Let 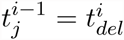and 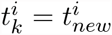, since the children of 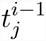 and 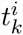 are the same, it is enough to consider *F*^*i-*1^. Then

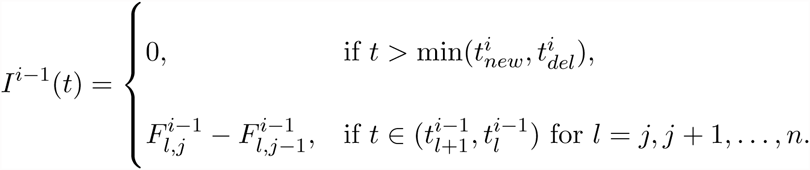

and

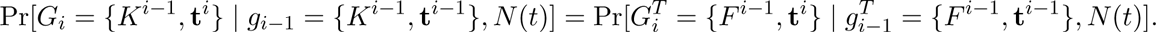

Now, when the deleted node corresponding to *t*_*del*_ is a cherry and the new node corresponding to *t*_*new*_ is also a cherry, there are four possible topologies *K*_*i*_ that lead to the same ranked tree shape *F ^i^*, then

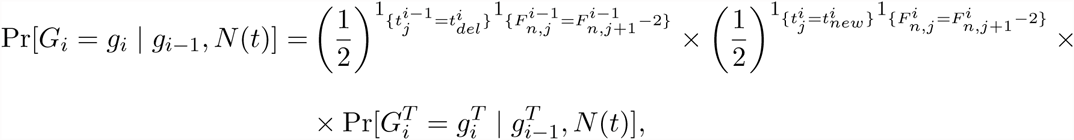

#### Proposition 5

*The marginal invisible transition density from a local Kingman’s genealogy g*_*i-*1_ *to G_i_ is equal to the marginal invisible transition density from the corresponding local Tajima’s genealogy* 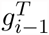 to 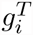.

*Proof.*

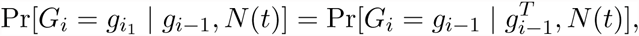

since all needed to compute the transition probability are the coalescent times and the *F*^*i-*1^ matrix. Since the topology does not change, the proof follows.

#### Proposition 6.

*The Likelihood of partially observed embedded SMC′ chain of local Kingman’s genealogies is proportional, up to a combinatorial factor, to the likelihood of partially observed embedded SMC′ chain of the corresponding local Tajima’s genealogies.*

*Proof.* The proof follows from propositions 3, 4 and 5 needed to express the likelihood of partially observed embedded SMC′ chain (Equation 13).

## Supporting Information

### 1 Visible Transitions

Figure S1A shows an example of a visible transition when the topology remains the same and Figure S1B shows an example of a visible transition when the topology changes. Green lines mark the possible pruning locations that could have lead to the same visible transition; the red circle indicates the deleted node at coalescent time *t*_*del*_ and the blue circle indicates the new node created at coalescent time *t*_*new*_.

**Figure S1:**
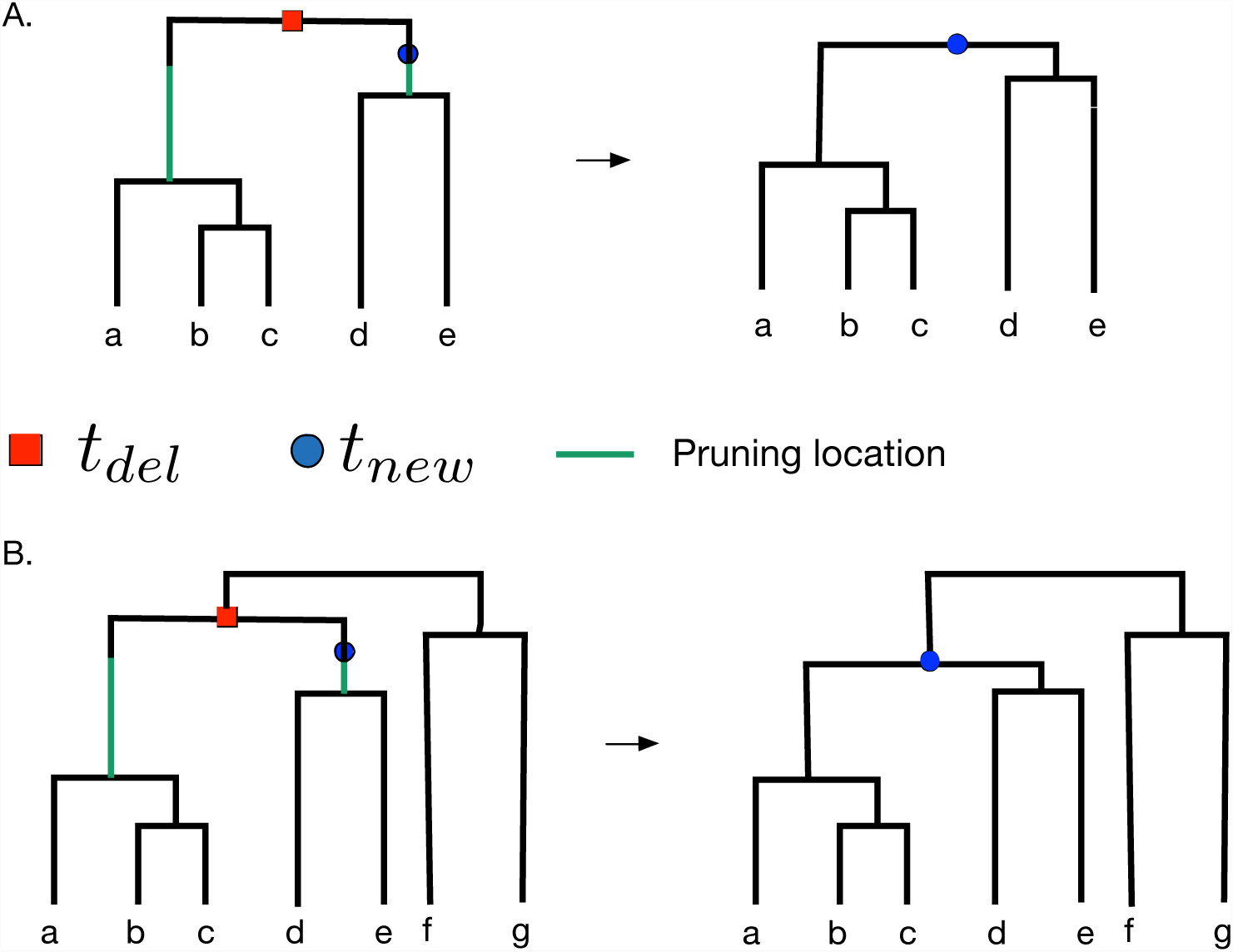
Examples of visible transitions when the pruning branch is uncertain. Red circle indicates deleted node at coalescent time *t*_*del*_, blue circle indicates new node at coalescent time *t*_*new*_. Green lines indicates possible pruning locations that could have resulted in such a visible transition. A. The topology remains the same. B. The topology changes.

**Figure S2:**
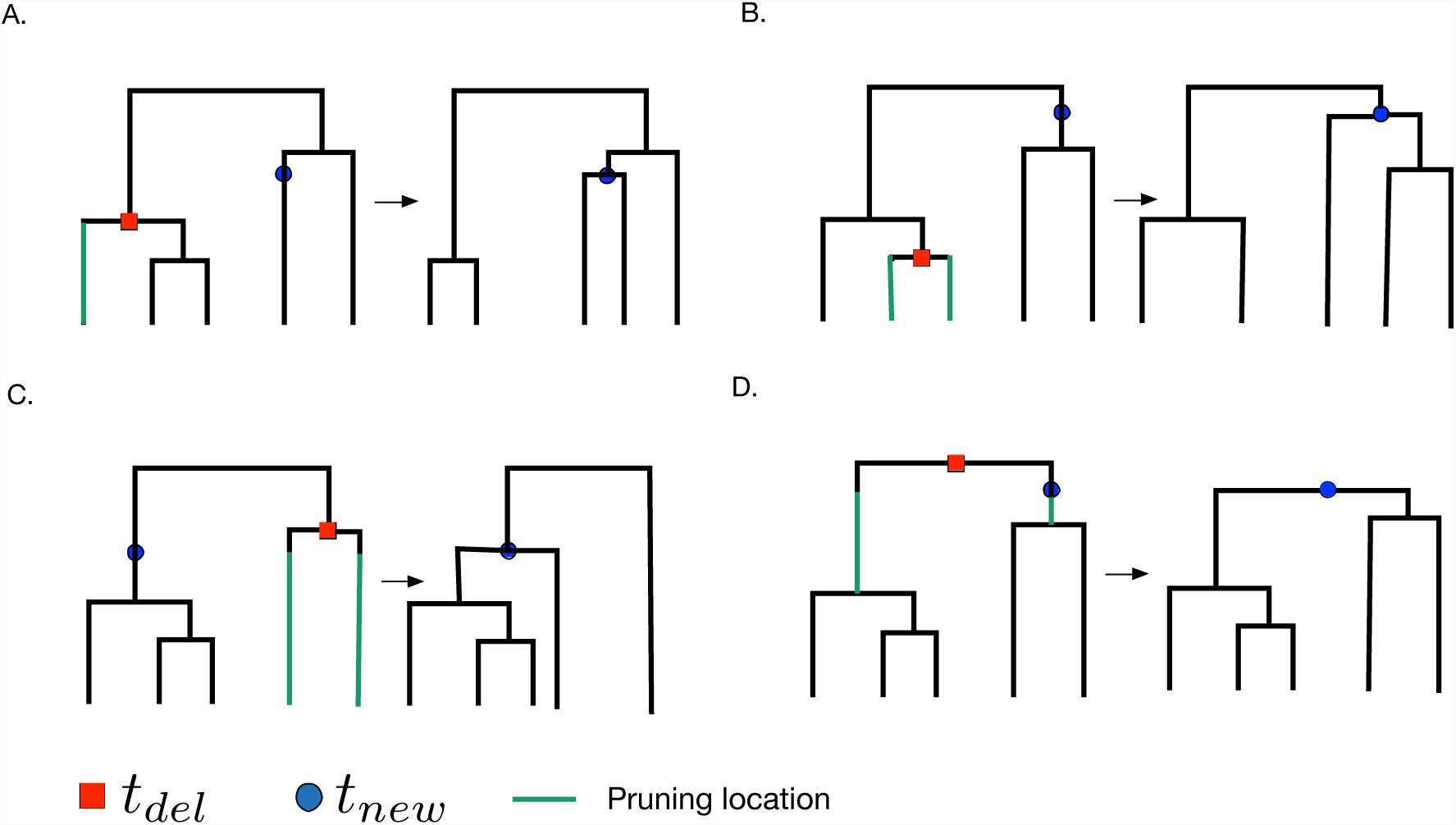
Examples of visible transitions between local Tajima’s genealogies. Red circle indicates deleted node at coalescent time *t*_*del*_, blue circle indicates new node at coalescent time *t*_*new*_. Green lines indicates possible pruning locations that could have resulted in such a visible transition.

### 2 Visible transitions between Tajima’s genealogies

A Tajima’s genealogy g*^T^* is an unlabeled genealogy. In Figure S2, we show four possible visible transitions. In the first case (Figure 2A), when we compare the number of *children* of the blue circle node on the right tree at time *t* with the *children* of the red circle node on the left tree, we can conclude that only the green branch could have been selected for pruning. In Figure 2B, comparing the *children* of the blue circle node on the right genealogy to the *children* of the red circle in the left genealogy, we conclude that the two *children* of the red circle are possible pruning locations. In Figures 2C-D, *t*_*new*_ < *t*_*del*_. This implies that the possible pruning locations will necessarily have heights up to *t*_*new*_. Again, by comparing the *children* of the blue circle node on the right to the *children* of the red circle node on the left, we can asses the possible pruning locations.

### 3 Simulations with MaCS

We use `MaCS` (Chen et al., 2009) for all our simulations with the following code lines:

Constant population size:

~~~
./macs2 300000 -t 1.0 -T -r .005 -h 1 (SEED: 1420480396)
./macs20 3000000 -t 1.0 -T -r .0002 -h 1 (SEED: 1399175725)
./macs100 3000000 -t 1.0 -T -r .0002 -h 1 (SEED: 1400528079)
~~~

Exponential growth and constant:

~~~
./macs2 300000 -t 1.0 -eG .1 10 -T -r .02 -h 1 (SEED: 1419985269)
./macs20 300000 -t 4.0 -eG .1 10 -T -r .002 -h 1 (SEED: 1420040333)
./macs100 300000 -t 1.0 -eG .1 10 -T -r .0002 -h 1 (SEED: 1401855826)
~~~

Bottleneck:

~~~
./macs2 300000 -t 4.0 -eN 0 1 -eN 0.3 0.1 -eN 0.5 1 -T -r .01 -h 1 (SEED: 1420824821)
./macs20 300000 -t 4.0 -eN 0 1 -eN 0.3 0.1 -eN 0.5 1 -T -r .002 -h 1 (SEED: 1420826310)
./macs100 300000 -t 4.0 -eN 0 1 -eN 0.3 0.1 -eN 0.5 1 -T -r .001 -h 1 (SEED: 1420826409)
~~~

**Figure S3:**
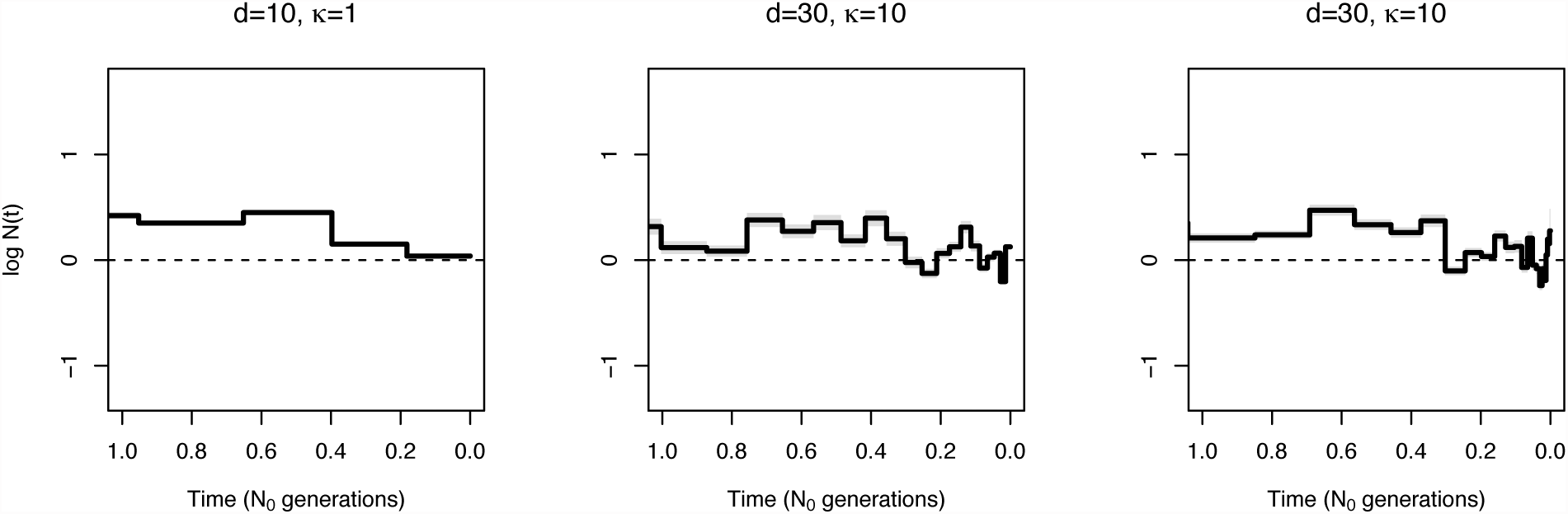
EM sensitivity to parameter discretization. Comparison of population size trajectories estimated from 1000 simulated genealogy (m = 1000) of 100 individuals with a constant population size. EM inference with different discretizations varying the parameters in Equation **??**.

**Table S1:** Summary of simulation results depicted in Figure S3. SRE is the sum of relative errors (Equation 24), MRW is the mean relative width of the 95% BCI (Equation 25), and ENV (Equation 26)

|  | SRE | MRW | ENV |
| --- | --- | --- | --- |
| EM $d = 10, \kappa = 10$ | 43.41 | 0.99 | 48.6% |
| EM $d = 30, \kappa = 10$ | 34.25 | 0.76 | 42.6% |
| EM $d = 30, \kappa = 100$ | 43.96 | 0.99 | 46.0% |

### 4 EM sensitivity to parameter discretization

In Figure S3, we show EM estimates of a constant population size from 1000 local genealogies of 100 individuals. We show that different discretizations result in different estimates. We note that confidence intervals perform poorly in terms of coverage. The performance statistics corresponding to the three estimations displayed in Figure S3 are shown in Table S1.

### 5 Analysis of Human data

We use *ARGweaver* (Rasmussen et al., 2014) with the following code lines:

European population:

~~~
arg-sample -s data1000/CEU_10.sites
   -N 11534 -r 1.6e-8 -m 1.26e-8
   --ntimes 200 --maxtime 200e3 -c 1 -n 10
   -o data1000/CEU.sample/out
~~~

Yoruban population:

~~~
arg-sample -s data1000/YRI_10.sites
   -N 11534 -r 1.6e-8 -m 1.26e-8
   --ntimes 200 --maxtime 200e3 -c 1 -n 10
   -o data1000/YRI.sample/out
~~~

**Figure S4:**
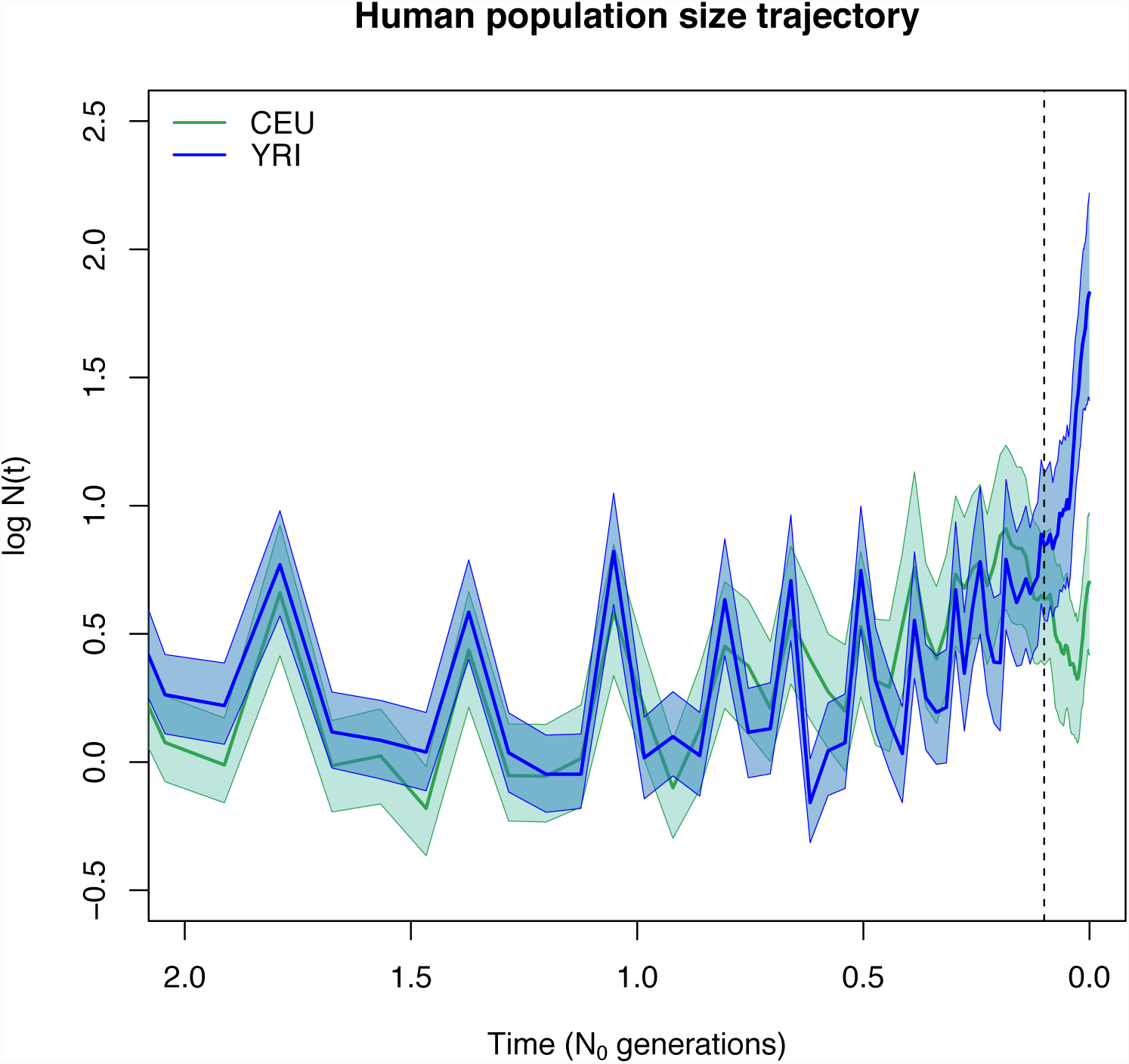
**Inference of human population size trajectories** *N*(*t*) **for *n*** = 10. Green solid line and green shaded areas represent the posterior median and 95% BCI for European population (CEU) and blue solid line and blue shaded areas represent the posterior median and 95% BCI for Yoruban population.

*ARGweaver* time is measured in units of generations, so in order to generate Figure 8, we multiplied time by 1/(2 × 11, 534). To obtain log *N*(*t*) displayed in Figure 8, we multiplied our estimates by 1/(8 × 11, 532) and converted them in logarithmic scale.

